# Nitrous oxide production, mechanisms, and modeling from a denitrifying phosphorus removal bioreactor

**DOI:** 10.1101/2024.09.04.611069

**Authors:** McKenna Farmer, Fabrizio Sabba, George Wells

## Abstract

Nitrous oxide (N_2_O) is a potent greenhouse gas produced as an unintentional, undesired byproduct in many nitrogen removal bioprocesses. Given the considerable challenges in managing N_2_O emissions from wastewater treatment, N_2_O could be reframed as a value-added product if intentionally generated and captured. This study assesses N_2_O production and mechanisms in a Coupled Aerobic-anoxic Nitrous Decomposition Operation with Phosphorus removal (CANDO+P) reactor. Optimal performance was achieved when the reactor was fed with a mixture of propionate and glucose, resulting in N_2_O was production up to 50% of influent nitrogen. Through 16S rRNA amplicon and shotgun metagenomic sequencing, we found that *Candidatus* Accumulibacter were the dominant phosphorus accumulating organism (PAO). Assembly of a high-quality metagenome-assembled genome showed that *Ca.* Accumulibacter encoded a full complete denitrification pathway from nitrite to nitrogen gas. We also found abundant populations of denitrifying glycogen accumulating organisms (GAO) and ordinary heterotrophic organisms (OHO). We also incorporated truncated denitrification pathways into a process model to predict N_2_O generation. N_2_O predictions were the most similar to observed results when the denitrification pathways of PAO, GAO, and OHO model populations reflected denitrification gene abundances from the metagenomic sequencing analysis. Our work demonstrates the feasibility of using non-VFA carbon for intentional N_2_O generation and provides broader insights into N_2_O generation and truncated denitrification pathways of denitrifying PAO and GAO.

## 1. Introduction

Nitrous oxide (N_2_O) is a greenhouse gas with approximately 300 times the global warming potential of carbon dioxide (CO_2_). N_2_O can constitute a majority of greenhouse gas emissions from wastewater treatment plants (Daelman et al., 2013), and N_2_O emissions from domestic wastewater treatment represent ∼5% of N_2_O emissions in the US (EPA, 2024). Reducing N_2_O emissions is an obvious target for reducing the carbon footprint of wastewater treatment, and significant research efforts are underway in pursuit of this goal. However, mitigating N_2_O emissions can be challenging due to competing operational goals. For example, reducing aeration is a common method to decrease energy consumption, thereby decreasing the carbon footprint of the process, but low dissolved oxygen conditions may lead to nitrifier-driven N_2_O production (Peng et al., 2015). Shortcut nitrogen removal methods such as nitritation-denitritation processes can also unintentionally increase N_2_O production due elevated nitrite (NO_2_^-^) concentrations (Zeng et al., 2003; Roots et al., 2020). Other factors may also increase N_2_O emissions besides the process configuration, including temperature (Hu et al., 2011), salinity (Shao et al., 2020), and influent carbon to nitrogen ratios (Velho et al., 2017). Given the challenges in managing N_2_O emissions from wastewater treatment, N_2_O could be reframed as a value-added product if intentionally generated and captured to prevent fugitive emissions.

N_2_O has a variety of uses and is a useful oxidant in combustion reactions. N_2_O has been used for decades in motorsports (Kelly, 2016) and has also been used as a propellant in aerospace applications (Gohardani et al., 2014; Werling et al., 2020). N_2_O is also being assessed for use as an oxidant in other applications, such as the production of functionalized phenols (Le Vaillant et al., 2022). Within a wastewater treatment facility, N_2_O gas could be used to increase power output from biogas combustion (Scherson et al., 2014). N_2_O gas is industrially produced by decomposing ammonium nitrate with heat, which presents significant environmental health and safety concerns. Furthermore, ammonium nitrate is derived from Haber-Bosch, an energy intensive process of nitrogen sequestration from the atmosphere that generates 1-2% of global CO_2_ emissions (Smith et al., 2020). Intentional N_2_O generation, capture, and combustion is an opportunity to redirect this potent greenhouse gas for a useful purpose.

Intentional N_2_O production from wastewater has been explored in the coupled aerobic−anoxic nitrous decomposition operation (CANDO) process (Scherson et al., 2014, 2013; Weißbach et al., 2018), which produces N_2_O through the denitrification pathway by feeding organic carbon and nitrogen (as NO_2_^-^) separately. The CANDO+P process (Gao et al., 2017) is a further development of the process where biological phosphorus removal is also incorporated through enrichment of denitrifying phosphorus accumulating organisms (DPAO). The CANDO+P process attempts to enrich a community where N_2_O reduction is minimized, while the preceding steps of the denitrification pathway are maintained such that N_2_O is the primary denitrification product. Like many other denitrifiers, DPAO and denitrifying glycogen accumulating organisms (DGAO) may possess incomplete denitrification pathways which are highly specific by species and strain (Albertsen et al., 2016; Petriglieri et al., 2022; Stewart et al., 2024). In general, the presence of NO_2_^-^ as an electron acceptor, alone or paired with other oxidized nitrogen species, increases N_2_O accumulation in DPAO and DGAO (Marques et al., 2018). Previous work in our lab on a CANDO+P system has found that different denitrifying populations in the same culture (DPAO, DGAO, and non-PAO/GAO denitrifiers) differentially expressed denitrification genes, resulting in a highly modular denitrification process where NO_2_^-^ reductases were also expressed at a higher level than N_2_O reductases (Wang et al., 2021). Similar results have been observed in a biological phosphorus removal system not intended for N_2_O production Vieira et al. (2018) as well as in other environmental contexts like soils (Bru et al., 2011; Nadeau et al., 2019).

The goals of this work are to (1) demonstrate CANDO+P for N_2_O accumulation on low COD:N wastewater with non-VFA carbon (2) understand distribution of denitrifying genes in the community and (3) predict N_2_O production through process models informed by molecular data. The results of this work are expected to increase our ability to optimize N_2_O accumulation by DPAO for resource recovery and can also be considered in the broader context of understanding N_2_O emissions from nutrient removal bioprocesses.

## 2. Results and Discussion

### 2.1. Reactor performance and N_2_O production on glucose-amended feed

The reactor was adapted for orthoP removal by approximately day 50 of operation (SI Figure 1). However, orthoP removal performance became unstable around day 110 and continued throughout the rest of the phase. This deterioration of performance seems to have coincided with changes in the microbial community, which is described further in section 2.2. Given the reactor instability, the COD:P ratio was increased to encourage PAO growth in Phase II. However, this change did not improve orthoP removal, and in a majority of sampling points, no net orthoP removal was achieved. NO_2_^-^ removal was consistent during this phase (average of 92 ± 16%), suggesting that conventional denitrifiers or DGAO were active during this phase rather than DPAO. During this phase, N_2_O accumulation was also observed during some anoxic phases but inconsistently and at low concentrations (<1 mgN/L).

Phase III (days 484-512) was used as a transition period to move the reactor from VFA-only to a mixed carbon feed. Previous work has demonstrated that PAO grown on acetate may have greater carbon uptake ability than GAO when the feed is switched to propionate (Oehmen et al., 2005), therefore the influent feed was amended with propionate on day 484 in a 1:1 COD ratio with acetate for a final concentration of 300 mg/L COD. During this phase, preferential uptake of propionate was observed, and acetate was carried over between cycles (SI Figure 2). NO_2_^-^ removal was consistent during this phase (average of 94 ± 10%), and N_2_O accumulation began to occur consistently, with peak values around 1 mgN/L, representing 5-10% conversion of influent NO_2_^-^-N.

Phase IV (days 512-750) was started by modifying the COD source to propionate and glucose on day 512 in a 2:1 COD ratio with an initial concentration of 250 mg/L COD. A typical in-cycle profile is shown in Figure 1. Unlike Phase III, there was no COD carry over between cycles, and COD was consumed completely within the anaerobic phase. During the initial part of the phase, there was a clear substrate preference for glucose over propionate (SI Figure 3). Rapid COD uptake was observed during this phase, so the anaerobic phase length was reduced from approximately 1.6 hours to 0.6 hours to prevent endogenous respiration, and the COD was reduced to 100 mg/L with a 1:1 propionate to glucose ratio on day 688. During Phase IV, orthoP removal was highly variable, reaching a maximum removal of 90%, but having a net negative average removal and large standard deviation (-34 ± 62%). In most cycles, we observed both anoxic and aerobic orthoP uptake (SI Figure 5). The mixed propionate and glucose feed resulted in NO_2_^-^ removal of 90 ± 14%, within the range of previous phases. As expected, the COD:N ratio had a strong, positive correlation with the anoxic orthoP uptake and NO_2_^-^ reduction rate (SI Figure 5). The ranges of anoxic orthoP uptake and NO_2_^-^ reduction were within ranges previously observed in a CANDO+P reactor (Gao et al., 2020). This phase had more consistent N_2_O accumulation than the previous phases, between 10-30% of the influent NO_2_^-^-N (SI Figure 4) and up to 50% during some cycles. The maximum specific N_2_O production rate was 0.96 ± 0.36 mgN_2_O-N/gTSS/h. Overall, the addition of glucose to the COD feed improved the consistency of N_2_O production but did not offer vast improvements to the orthoP removal performance.

**Figure 1.**
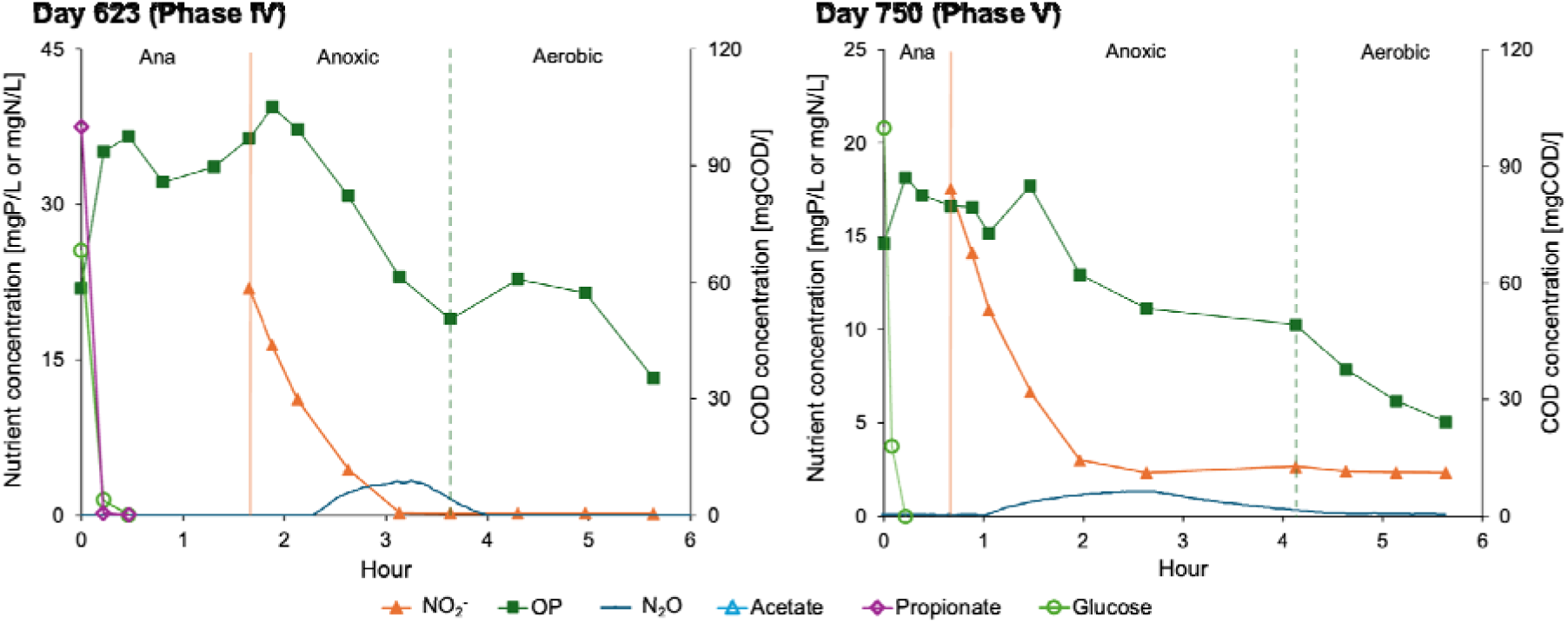
Representative in-cycle nutrient and carbon profile from days 623 and 720.

In the final operational phase (Phase V), the COD feed was switched to glucose only to test the viability of CANDO+P on non-VFA carbon. On the first day of glucose-only operation (day 720), denitrification was not complete (Figure 1), suggesting that the propionate dosed in Phase IV provided important benefits to carbon storage products used for denitrification. In this phase, we observed a shift from unstable phosphorus removal to more robust performance. In the first month of the phase, the effluent orthoP was the same as or higher than the influent, and the orthoP removal averaged over the whole phase was highly variable (average of 32 ± 54%). However, towards the end of the phase, effluent orthoP reached the detection limit (0.02 mgP/L, ∼99% removal efficiency). N_2_O accumulation occurred during this phase but was lower in magnitude and rate than Phase IV. Influent NO_2_-N was gradually increased up to 30 mgN/L in this phase, but peak N_2_O values were between 1-2 mgN/L, representing only up to 6% conversion of influent NO_2_-N, with a maximum observed N_2_O production rate of 0.25 mgN_2_O-N/gVSS/h. Although the glucose-only feed provided an improvement to orthoP removal performance, this COD source resulted in lower N_2_O production on its own compared to the previous phase with VFA and glucose. While detrimental for our goal of N_2_O production, this result has positive implications for processes fed with significant quantities of glucose or possibly other simple sugars where N_2_O production is undesirable.

While the CANDO+P reactor in this work achieved consistent N_2_O production in Phases IV and V with glucose as a COD source, the N_2_O production was lower than previous studies. For example, Gao et al. (2017) achieved an N_2_O conversion efficiency of 70% of influent NO_2_^-^-N compared to up to the typical value of 30% and maximum value of 50% achieved in this work. The N_2_O production rate of 0.96 ± 0.36 mgN_2_O-N/gVSS/h in Phase IV was also much lower than previous values on acetate-only (5.7 ± 2.0 mgN_2_O−N/gVSS/h) and propionate-only (4.9 ± 2.1 mgN_2_O−N/gVSS/h) observed by Gao et al. (2017). However, the N_2_O production rate observed here was consistent with observations from a pilot fed with real wastewater with a rate of 1.30 ± 0.20 mgN_2_O−N/gVSS/h (Weißbach et al., 2018). A CANDO reactor was also recently operated with conversions near 100% in some cycles (Wang et al., 2020), much higher than achieved here, but Wang and colleagues also observed cycles where complete N_2_O was produced and completely reduced before the end of the anoxic phase, similar to what we observed in Phase IV and V of this study. To better understand the microbial drivers of N_2_O production in this reactor, we next turned to a suite of complementary molecular microbial community analyses.

### 2.2. Denitrifying community shifts over the reactor operation

16S rRNA amplicon sequencing was used to identify prevalent PAO, GAO, and denitrifier taxa in the reactor. Relative abundances from 16S rRNA sequencing for the whole study are shown in Figure 2. As previously mentioned, orthoP removal performance in Phase I was unstable after an increase in the NO_2_^-^ dose. The only PAO detected in Phase I, *Ca*. Accumulibacter, increased in abundance until approximately day 125 but tapered off and eventually declined over the rest of the phase. On the other hand, the relative abundance of the GAO *Plasticicumulans* increased, reaching its peak around day 150, when orthoP instability was greatest. *Plasticicumulans* are in the *Competibacteraceae* family and are generally considered GAO due to their uptake of soluble COD under feast-famine conditions (McIlroy et al., 2014). These organisms do not have denitrification capabilities (Tamis et al., 2014), so denitrification was likely carried out by other organisms during this unstable period, potentially *Dechloromonas* which increased in abundance after day 125. Some species of *Dechloromonas* possess the polyphosphate accumulating metabolism, but these species were not detected in Phase I. Another denitrifier present during Phase I was *Flavobacterium,* which have previously been associated with other nitrite-fed phosphorus removal bioreactors (Lv et al., 2014; Wang et al., 2024). Overall, the decline of *Ca.* Accumulibacter PAO and concurrent GAO and denitrifier community shifts likely contributed to the poor orthoP removal performance during Phase I.

**Figure 2.**
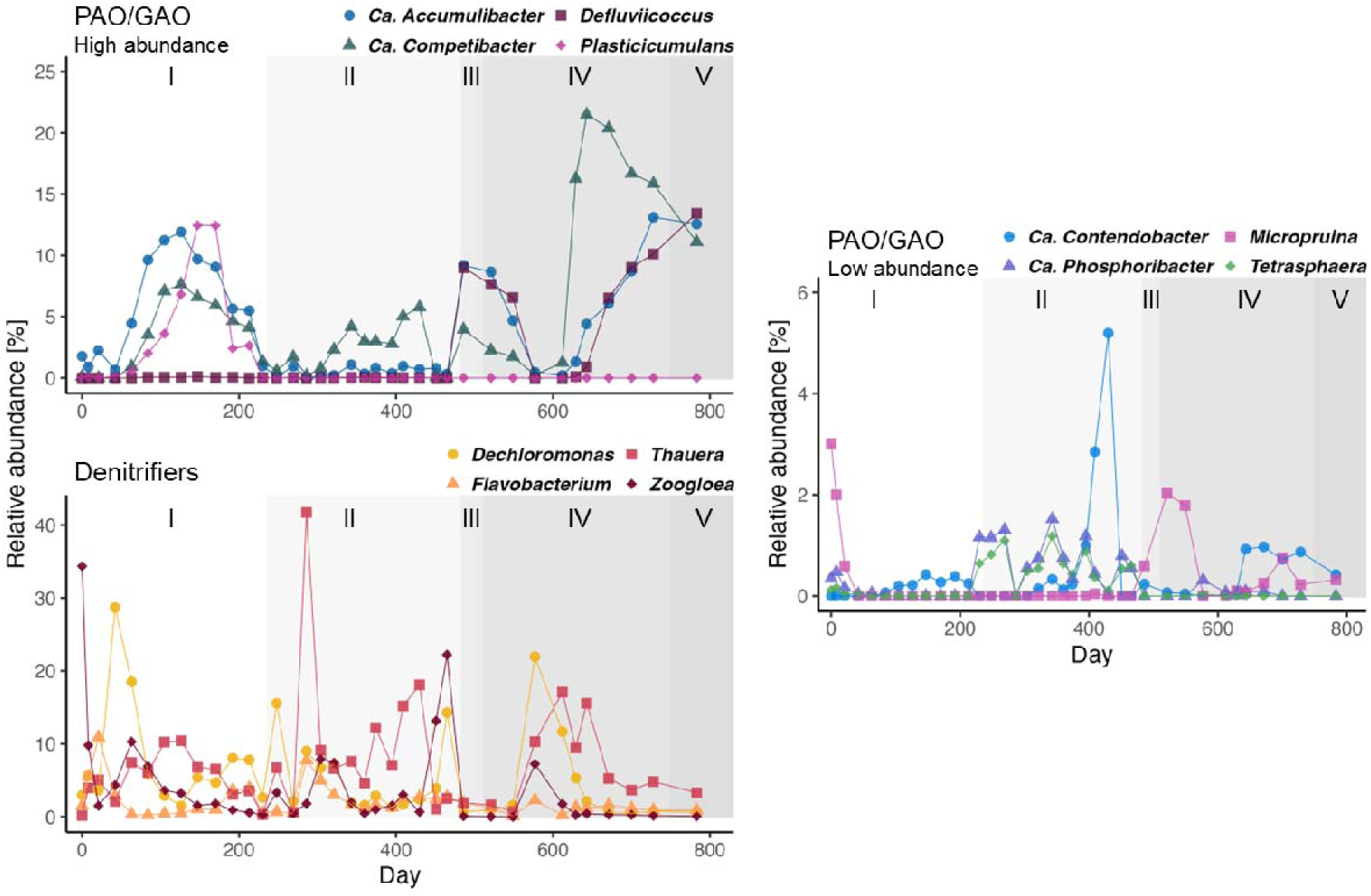
PAO, GAO, and denitrifier relative abundances from 16S rRNA sequencing. Phases (I-V) are denoted by the shaded areas.

During Phase II (acetate and high NO_2_^-^ feed), which had largely unstable orthoP removal, *Ca.* Accumulibacter abundance dropped to a median of 0.4% compared to 5.5% in Phase I. In place of *Ca.* Accumulibacter, the PAO *Ca.* Phosphoribacter and *Tetrasphaera* increased in relative abundance, though not to the same magnitude as *Ca.* Accumulibacter (Figure 2). These organisms were present in the seed sludge but had been at negligible abundances throughout Phase I. Similarly, the GAO *Ca.* Contendobacter was present during Phase I but had a greater abundance in Phase II and a notable spike around day 400. The GAO *Plasticicumulans* did not reemerge in this phase or subsequent phases.

The transitional Phase III (acetate and propionate feed) corresponded to an increase in *Ca.* Accumulibacter, as well as the GAO *Ca.* Competibacter and *Defluviicoccus.* The sudden enrichment of *Defluviicoccus* may have been related to the addition of propionate, as previous work has demonstrated that *Defluviicoccus* will uptake propionate even in the presence of other substrates like acetate (Burow et al., 2007). *Defluviicoccus* has also been observed at high abundances in other denitrification systems with significant NO_2_^-^ concentrations, including a CANDO system (Wang et al., 2020) and a partial-denitrification anammox system (Chu et al., 2021), though other works show *Defluviicoccus* suppressed in the presence of NO_2_^-^ (Tayà et al., 2013).

In Phase IV (propionate and glucose feed), PAO and GAO abundances decreased initially, then rebounded after approximately 60 days. Notably, we documented a strong positive correlation between *Ca.* Accumulibacter and *Defluviicoccus*, which had a Pearson correlation of 0.94 (p < 0.001). A correlation is reasonable given the suitability of feast-famine feeding for both of these organisms, but the strength of the correlation and nearly identical abundances was surprising. Another notable feature was the rapid increase in *Ca.* Competibacter in Phase IV, which was driven by two strains of *Ca.* Competibacter that were not highly abundant prior to day 600 (SI Figure 6). Additionally, the GAO *Micropruina* also increased in abundance at the beginning of the phase. *Micropruina* can directly uptake glucose under anaerobic conditions (McIlroy et al., 2018), giving them a competitive advantage for carbon uptake in this phase. Overall, the net abundances of GAO were much greater than PAO in Phase IV and V, which may have contributed to the inconsistent orthoP removal performance observed in this time period.

### 2.3. Microbial community features driving N_2_O accumulation

Shotgun metagenomic sequencing was used to understand denitrifying capabilities of the microbial community. Metagenomic samples were collected primarily during Phase IV. In this phase, a mix of propionate and glucose were added to the feed, and N_2_O was produced then mostly reduced by the end of the anoxic phase. The sequencing data was first analyzed using a pathway-focused metagenomics approach for denitrification pathway genes by annotating coding sequences from contigs. Figure 3 summarizes taxa relative abundances and *nosZ* gene abundances during a period of Phase IV when N_2_O production decreased. A notable feature is the prevalence of *nosZ* associated with *Ca.* Accumulibacter, while *nosZ* associated with other taxa decreased. *Thauera* and *Ca.* Competibacter abundances increased based on 16S rRNA sequencing, but there was a net decrease in *Thauera-*associated *nosZ* and no increase in *Ca*. Competibacter-associated *nosZ* over the same time period. These results suggest that *Ca.* Accumulibacter contributed to N_2_O reduction more so than other community members, though we did not measure gene expression so cannot say definitively.

**Figure 3.**
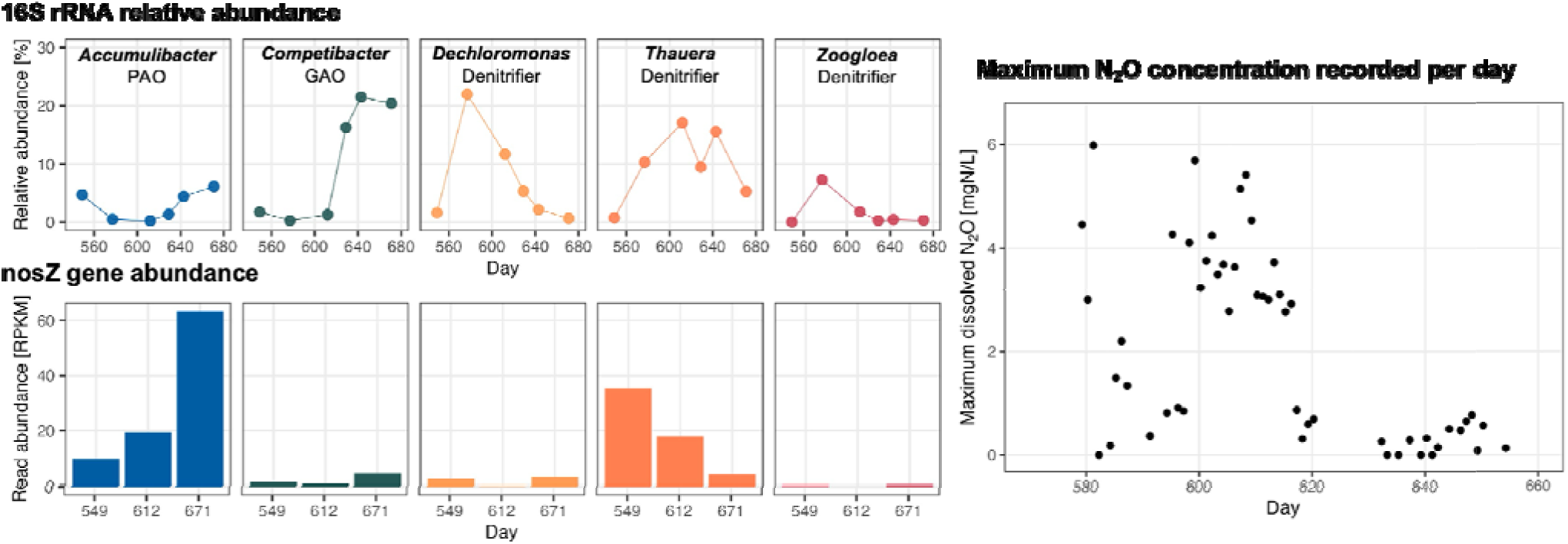
Relative abundances of key PAO, GAO, and denitrifier taxa from 16S rRNA sequencing (top left), *nosZ* gene abundance from metagenomic sequencing (bottom left), and maximum N_2_O concentrations recorded during part of Phase IV (right).

NO_2_^-^ reduction was a common function across almost all analyzed taxa (SI Figure 7), though with some variances in the specific corresponding functional gene. The genes *nirA* and *nirBD* encode assimilatory NO_2_^-^ reduction (i.e. NO_2_^-^ reduced to NH_3_ for incorporation into biomass) while *nirS* and *nirK* encode dissimilatory NO_2_^-^ reduction (i.e. respiration). *Defluviicoccus* was the exception to this, as the only *Defluviicoccus-*associated denitrification genes detected (*norBC*) encoded NO reduction and were at relatively low abundances compared to those from other organisms. This finding suggests that *Defluviicoccus*, though a dominant GAO by abundance, were not significant denitrifiers in this system. Other *norBC* genes were identified from the GAO *Ca.* Competibacter and *Ca.* Contendobacter as well as the denitrifiers *Dechloromonas, Thauera,* and *Zoogloea*. *Ca.* Accumulibacter-associated *norBC* genes increased in abundance over Phase IV unlike the denitrifier abundance, which decreased to undetectable levels by the end of the phase. In this reactor, *Ca.* Accumulibacter putatively encoded genomic capability for denitrification from NO_2_^-^ to N_2_. This metagenomic analysis cannot be used to determine if the genes were all present in one organism, so a binning approach was undertaken to recover more information about *Ca.* Accumulibacter and other denitrifying community members.

We employed a coupled assembly and binning approach to recover metagenome-assembled genomes (MAGs). Medium-(≥50% completeness, ≤10% contamination) and high-quality (≥90% completeness, ≤5% contamination) PAO, GAO, and denitrifier MAGs are summarized in SI Table 1, and associated denitrification genes are shown in SI Figure 8. Two *Ca.* Accumulibacter MAGs were recovered: one high-quality (CAN3.12, completeness 97.95%, contamination 1.98%) and one medium-quality (CAN_3.27, completeness 74.14%, contamination 1.72%). The *Ca.* Accumulibacter MAGs possessed genes for a complete denitrification pathway from NO_2_^-^ to N_2_, which is in agreement with the denitrifying pathway metagenomics results. However, the NO reductase genes present in the assembled MAGs (*norV*) differed from the dominant genes in the pathway approach (*norB* and *norC*). Some *norVW* genes associated with *Ca.* Accumulibacter were detected with the pathway approach but at very low abundances (<1 RPKM).

Both *Ca.* Accumulibacter MAGs possessed *glk* genes, which encode glucokinase, an enzyme that phosphorylates glucose to glucose-6-phosphate. This molecule is an entry point for numerous cellular functions, including glycolysis and glycogen synthesis. The presence of *glk* in the *Ca.* Accumulibacter MAGs suggests that these organisms have a direct pathway for glucose utilization. Our findings mirror a recent study which reported the direct uptake and utilization of glucose by *Ca.* Accumulibacter as well as assembly of a *Ca.* Accumulibacter MAG with the *glk* gene (Ziliani et al., 2023). It is unclear if *Ca.* Accumulibacter use glucose directly in glycogen storage, or if they can ferment glucose to pyruvate, then use this for PHA storage. In this study, we observed what appears to be a two phase orthoP release in Phase IV when propionate and glucose were dosed together, despite both carbon sources being taken up at a similar rapid rate (Figure 1). This work cannot be used to draw any definitive conclusions on the specific glucose utilization pathways by *Ca.* Accumulibacter but does show a potential for non-VFA carbon uptake and orthoP release by this canonical PAO.

### 2.4. Microbiome-informed mechanistic model

A mechanistic mathematical process model was developed to simulate N_2_O production during Phase IV using three denitrifying populations: DPAO, DGAO, and denitrifying OHO with internally stored carbon. Three scenarios were evaluated based on the denitrification pathway metagenomics results. The “full pathways” scenario was evaluated assuming DPAO, DGAO, and OHO performed complete reduction of NO_2_^-^ to NO to N_2_O to N_2_. The “truncated DGAO” scenario was evaluated assuming complete denitrification by DPAO and OHO, and no N_2_O reduction by DGAO based on the low DGAO-associated *nosZ* gene abundances observed during Phase IV (Figure 3, SI Figure 7). The “truncated DGAO+OHO” scenario was evaluated assuming complete denitrification by DPAO, and truncated denitrification pathways in both DGAO and OHO based on low *nosZ* and *norBC* gene abundances in these functional groups (**Error! Reference source not found.**). Rate equations, stoichiometry, and parameter values were based on the work of <u>Liu et al. (2015)</u> for DPAO, <u>Ren et al. (2023)</u> for DGAO, and <u>Gujer et al. (1999)</u> for OHO, and further details are provided in the supporting material.

N_2_O profile predictions compared to the observed values on day 623 are shown in Figure 4. The full pathways scenario did not predict any significant N_2_O accumulation, deviating the most from the observed values, and had a root mean square error (RMSE) value of 2.21. The truncated DGAO scenario predicted some N_2_O accumulation but was not consistent with the leading edge of the N_2_O profile or the overall magnitude and had an RMSE value of 1.36. The truncated DGAO+OHO scenario had the best match to the observed N_2_O profile in both shape and magnitude with an RMSE value of 0.68. Similar to the N_2_O profile, the NO_2_^-^ profile (SI Figure 10) was best captured by the truncated DGAO+OHO scenario (RMSE = 1.12), followed by the truncated DGAO (RMSE = 1.68) and full pathway (RMSE = 2.49) scenarios.

**Figure 4.**
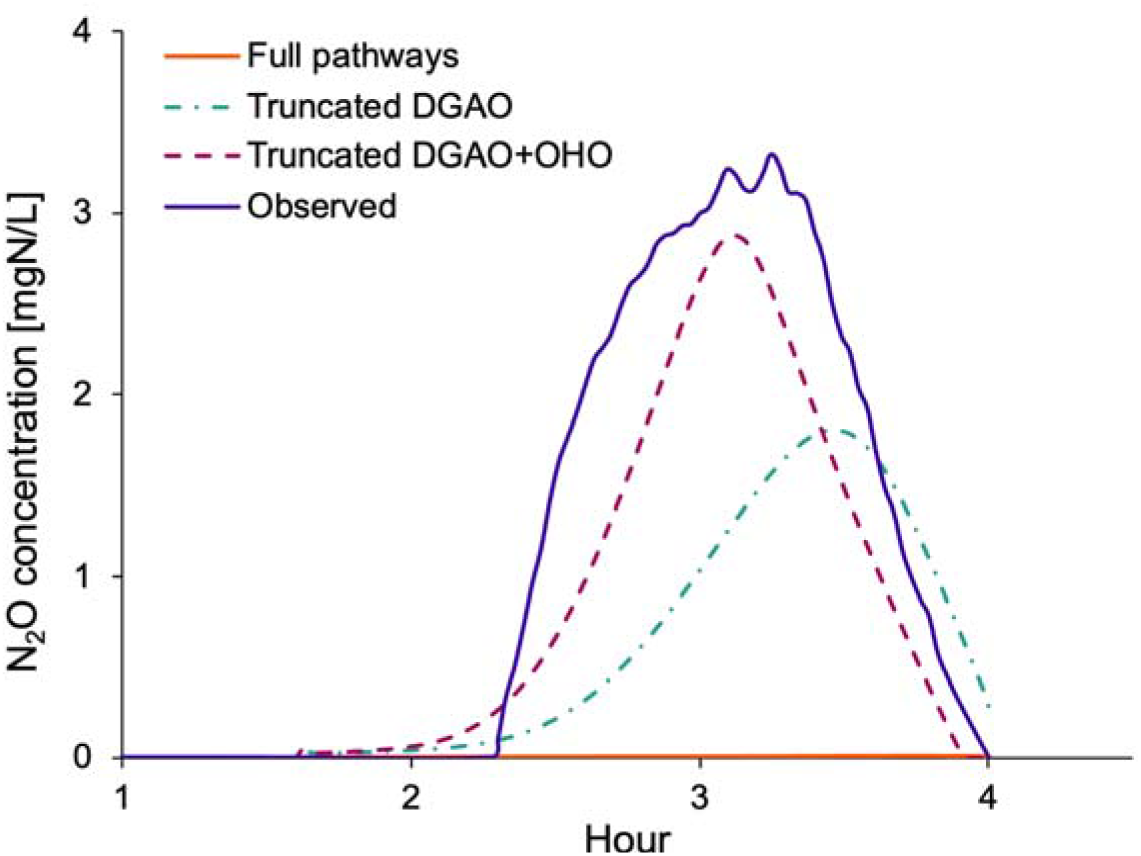
Predicted N_2_O profiles compared to the observed profile from day 623.

Future iterations of this model can improve the accuracy of the N_2_O and NO_2_^-^ profiles, potentially through incorporation of more accurate inhibition effects from NO_2_^-^. Ribera-Guardia et al. (2016) demonstrated that DGAO-enriched cultures produced more N_2_O than DPAO-enriched cultures when fed NO_2_^-^, likely due to a greater inhibitory effect of NO_2_^-^ on DGAO than DPAO. The importance of NO_2_^-^ inhibition factors is demonstrated in this work in SI Figure 11, where the truncated DGAO+OHO scenario was run assuming the same inhibition constant (8 gN/m^3^) for all denitrification steps. In this scenario, the peak N_2_O predicted was 4.2 mgN/L compared to 2.9 mgN/L in the original model, an increase of around 50%. While this model could be improved through more rigorous calibration, particularly for NO_2_ inhibition, the results demonstrate the utility of incorporating full and partial denitrification pathways by multiple functional groups to improve N_2_O predictions.

We also assessed the accuracy of the model in fitting the observed orthoP profile (SI Figure 10). All three modeling scenarios largely captured the magnitude of orthoP release and uptake. The two phase orthoP release observed in Phase IV was not captured by the model. This was unsurprising as we used one state variable for soluble carbon (S_S_) rather than individual variables for glucose and propionate. Existing model structures partition carbon into different categories to capture carbon availability dynamics over time, such as ASM2d which includes fermentation products (S_A_), fermentable readily biodegradable substrate (S_F_), and slowly biodegradable solids (X_S_) (Henze et al., 1999). This type of model structure can serve as an example of carbon partitioning, though in this work, it is obvious that both VFA and glucose are readily biodegradable and suitable for rapid uptake. Despite this limitation of our model on carbon uptake dynamics, all three scenarios performed reasonably well at predicting net orthoP release and anoxic orthoP uptake.

Overall, our predictions for N_2_O, NO_2_^-^, and orthoP relative to the observed data provide compelling evidence that incorporating truncated denitrification pathways into process models based on metagenomics data can be a useful route to predicting N_2_O production and consumption in DPAO and DGAO enriched bioprocesses. The goal of this modeling work was not to precisely calibrate and validate the model but rather to demonstrate the utility of this approach. Metagenomics-informed modeling has been tested with varying levels of success in soils (Graham et al., 2016; Nadeau et al., 2019) and similar approaches have been tested more recently in wastewater process modeling specifically, including modeling simplified partial denitrification (NO_3_^-^ to NO_2_^-^, NO_2_^-^ to N_2_) of heterotrophic denitrifiers (Su et al., 2023) and nitrite oxidizing bacteria (NOB) (Sampara et al., 2022). Metagenomics-informed modeling has even been tested in a denitrifying EBPR process with mixed populations of DPAO and DGAO (Oehmen et al., 2010). The approach presented here builds upon the work of Oehmen et al. (2010) by developing the model structure from metagenomic data collected from the reactor of interest, rather than relying on metagenomic data published from other study systems. This metagenomics-informed modeling approach would need further validation for full-scale activated sludge. More complicated microbial communities and influent immigration of microbes from the conveyance system would likely dilute the signal of denitrification genes. Furthermore, there are a variety of factors besides truncated denitrification pathways in heterotrophs which impact N_2_O production, particularly nitrifier-derived N_2_O production, oxygen inhibition, and transitional conditions between redox zones (Vasilaki et al., 2019).

## 3. Conclusion

A CANDO+P reactor was operated for over two years and evaluated for performance on VFA and non-VFA carbon, denitrifying microbial community dynamics, and utility of metagenomics data for improving N_2_O modeling. The CANDO+P reactor had unstable performance on an acetate-only feed, possibly due to an overabundance of the GAO *Plasticiculumans*. N_2_O production rate and reliability improved when the propionate feed was supplemented with glucose in Phase IV, reaching up to 50% conversion of influent nitrogen with a production rate of 0.96 ± 0.36 mgN_2_O-N/gVSS/h. This production was significantly higher than typical denitrification processes but fell short of previous CANDO/CANDO+P studies. Microbial community analysis via 16S rRNA sequencing showed that *Ca.* Accumulibacter were the dominant PAO throughout the study, and while GAO *Ca.* Competibacter and *Defluviicoccus* were also persistent throughout the reactor operation. Denitrification pathway analysis via shotgun metagenomic sequencing provided evidence that *Ca.* Accumulibacter populations may have encoded a complete denitrification pathway from NO_2_^-^ to N_2_ while GAO had low and nondetectable abundances of *nosZ*. A denitrification model with three denitrifying populations (DPAO, DGAO, OHO) was developed that incorporated truncated denitrification pathways based on the metagenomic sequencing data. The models with truncated denitrification pathways informed by metagenomic sequencing improved N_2_O predictions over scenarios assuming full denitrification pathways. This work demonstrates combined N_2_O production and orthoP removal through the CANDO+P process on non-VFA carbon, provides broader insights into truncated denitrification pathways of denitrifying PAO and GAO, and proposes metagenomic-informed modeling to improve predictions of N_2_O generation.

## 4. Materials and Methods

### 4.1. Reactor operation and conditions

A 12 L reactor was seeded with biomass from Stickney Water Reclamation Plant in May 2021 and operated for 830 days. The reactor was operated to select for DPAO through the primary react phases: fill and anaerobic react, nitrite (NO_2_^-^) dose and anoxic react, and aerobic react followed by settling and decanting. NO_2_^-^ was used as the nitrogen source to simulate effluent from an upstream nitritation reactor. The SBR was controlled using ChronTrol XT 4 circuit programmable timers. Complete SBR cycles were 8 hours, and the reactor decanted 6 L each cycle. Synthetic wastewater was used in this work and is detailed in the supplementary information. The phases of reactor operation were based on the COD source, COD concentration, and NO_2_^-^ dose (Table 4-1). Phase I was designated as the start-up phase but was operated longer than expected due to performance upsets. The mixed liquor suspended solids (MLSS) was an average of 0.76 ± 0.30 g/L during Phases I and II and increased to an average of 5.1 ± 2.1 g/L from Phase III until the end of the study.

**Table 4-1.**
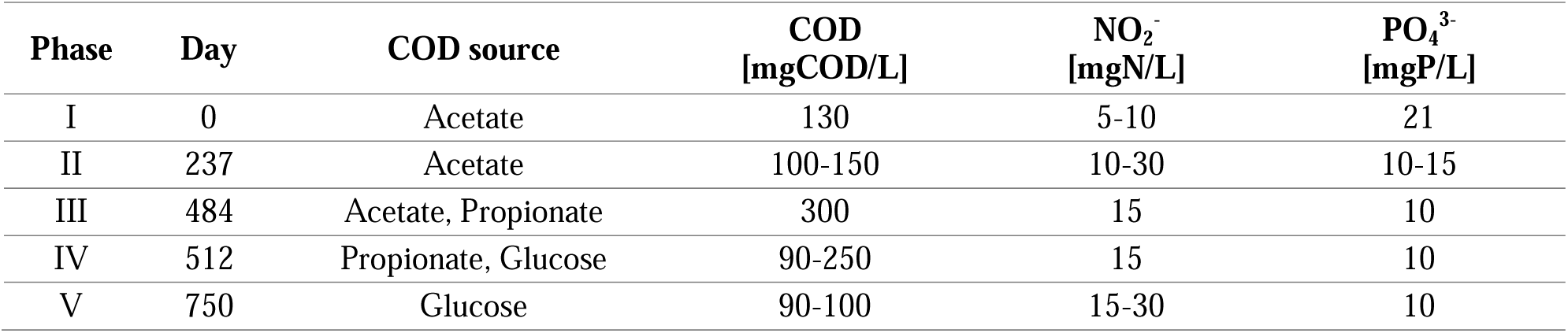
Reactor operational phases.

### 4.2. Water quality and reactor performance measurements

Reactor effluent samples were collected 2-3 times per week and measured for total and soluble chemical oxygen demand (COD), orthoP, NO_x_ (NO_3_^-^ + NO_2_^-^), and NO_2_^-^ per APHA standard methods (APHA, 2017). Nutrient measurements were conducted on a Skalar San++ continuous flow analyzer. Reactor solids samples were collected 1-2 times per week for total suspended solids (TSS) and volatile suspended solids (VSS) measurement per APHA standard methods. Dissolved N_2_O concentrations were monitored with a Unisense N_2_O Wastewater System. High time resolution in-cycle tests were conducted approximately every week to measure soluble orthoP, NO_2_^-^, volatile fatty acids (VFA), and glucose over the course of a reactor cycle. VFA measurements for acetate and propionate were performed on a GC-FID following APHA standard methods. Soluble glucose concentrations were measured with Sigma-Aldrich and Thermo-Fisher hexokinase measurement kits following manufacturer protocols.

### 4.3. Molecular sampling and DNA sequencing

Biomass was archived weekly in 3-4 replicates for DNA-based analysis. 1.5 mL aliquots of biomass were centrifuged at 10,000G for 3 minutes and decanted, and samples were resuspended with 1 mL of tris-EDTA buffer. The samples were centrifuged and decanted again, then pellets were stored at -80°C until further processing. DNA was extracted with the FastDNA SPIN Kit for Soil (MP Biomedicals) following the manufacturer protocol.

PCR amplification was performed on extracted DNA to amplify the V4-V5 region of the 16S rRNA gene using 515F/926R primers (Parada et al., 2016). PCR details are in the supporting information. PCR products were sent to Rush University Genomics and Microbiome Core Facility (Chicago, IL) for barcoding via second stage PCR and 2×300 bp sequencing with an Illumina MiSeq using V3 chemistry. Raw sequencing reads were processed using QIIME2, and amplicon sequence variants (ASV) were produced using deblur2. Taxonomy was assigned using the MiDAS v5.3 database (Dueholm et al., 2024).

Extracted DNA from Phases III and IV (days 451, 521, 549, 612, 671, 728) was sent to either Northwestern University NUSeq Core or SeqCenter (Pittsburgh, PA) for shotgun metagenomic sequencing. Library preparation was performed at both facilities with the Illumina DNA Prep kits following manufacturer protocols. Sequencing at the NUSeq Core was performed on a NovaSeq 6000 to generate 2×150 bp reads. Sequencing at SeqCenter was performed on a NovaSeq X Plus to generate 2×151 bp reads. Metagenomic analysis was performed on the Quest High Performance Computing Cluster at Northwestern University. Denitrification pathway analysis was performed by read trimming with fastp, assembly with MEGAHIT, prediction of protein-coding genes with Prodigal, and alignment of coding sequences to denitrification genes from UniProtKB (accessed May 2024). Coding sequences were retained with 70% or greater sequence identity to the reference. Metagenome-assembled genome analysis was performed on the assembled contigs from MEGAHIT, which were binned using metabat2, assigned taxonomy using GTDB-Tk, and annotated with prokka. Analysis scripts are available at https://github.com/mckfarm/cando_meta. All raw reads are available at accession PRJNA1148969.

### 4.4. Modeling N_2_O dynamics

Process modeling was used to understand whether predictions of N_2_O generation could be improved with denitrification pathway information from metagenomic sequencing. Three populations were used in the model: DPAO, DGAO, and denitrifying ordinary heterotrophic organisms (OHO). The stoichiometry, rate equations, and parameters were based on Liu et al. (2015) and Ren et al. (2023), for DPAO and DGAO, and ASM3 (Gujer et al., 1999) and Liu et al. (2015) for OHO. The reactor was modeled in AQUASIM as a single completely mixed compartment (Reichert, 1994). Each denitrifying population was modeled to perform anaerobic carbon storage, and the default modeling scenario included denitrification on NO_2_^-^, NO, and N_2_O for all three populations. Denitrification pathway distribution of the three populations from metagenomic data was used to generate multiple test scenarios of truncated denitrification pathways in DGAO and OHO. The stoichiometric matrix, rate equations, and parameter values are detailed in the supporting information.

## 5. Acknowledgements

McKenna Farmer and George Wells were supported in part by the Israel-U.S. Collaborative Water-Energy Research Center (CoWERC), via the Binational Industrial Research and Development Foundation (BIRD) Energy Center grant EC-15. This research was also supported in part through the computational resources and staff contributions provided for the Quest high performance computing facility at Northwestern University which is jointly supported by the Office of the Provost, the Office for Research, and Northwestern University Information Technology.

