## Supplemental Information for "Nitrous oxide production, mechanisms, and modeling from a denitrifying phosphorus removal bioreactor"

### Synthetic wastewater media

The synthetic wastewater mineral media contained the following: 70 mg/L MgSO_4_, 20 mg/L CaCl_2_, 1 mg/L yeast extract, and 100 mg/L NaHCO_3_. Each liter of media also contained 0.3 mL of Hutner’s trace elements solution, consisting of 5 g/L FeSO_4_·7H_2_O, 50 g/L EDTA, 22 g/L ZnSO_4_·7H_2_O, 11 g/L H_3_BO_3_, 5 g/L MnCl_2_·4H_2_O, 1.6 g/L CoCl_2_·6H_2_O, 1.6 g/L CuSO_4_·5H_2_O, and 1.1 g/L of (NH_4_)_6_Mo_7_O_24_·4H_2_O, with KOH pellets added to adjust the pH to 6.5. COD and P media was prepared with NaAc, NaPr, and/or (D)-glucose, as well as equal parts NaH_2_PO_4_·H2O and K_2_HPO_4_ on a phosphorus mass basis. NO_2_^-^ media was prepared with NaNO_2_.

### PCR information

Each PCR reaction used 20 uL reaction volumes containing 10 uL DreamTaq Green PCR Master Mix (Thermo Scientific, Waltham, MA, USA), 0.5 uM each of forward and reverse primer (IDT, Coralville, IA, USA), 2 uL of template DNA, and molecular grade water. The following thermocycling conditions were used as described in Parada et al. (2016) and optimized for DreamTaq Green: 95°C for 3 min; 25 cycles of 95°C for 45 seconds, 50°C for 60 seconds, and 68°C for 90 seconds; 68°C for 7 minutes. The forward and reverse primers were appended with Fluidigm Access Array linkers. PCR product length was confirmed using gel electrophoresis.

### Supporting Figures

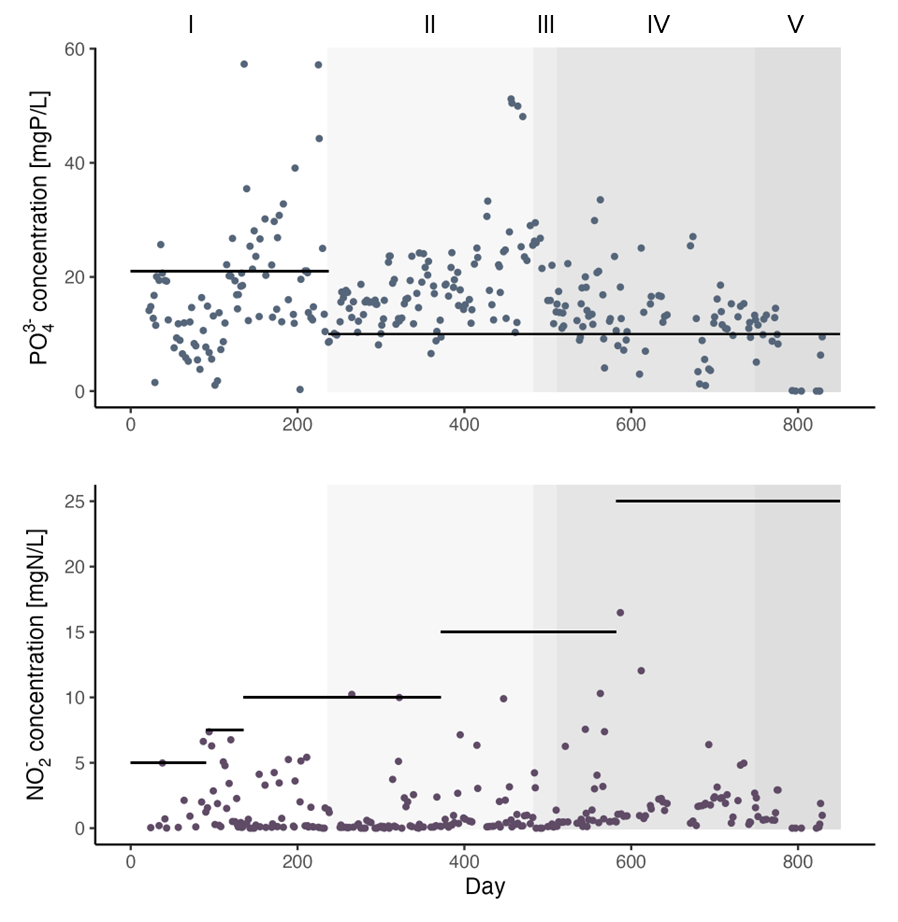

Figure 1 Effluent orthoP (top) and nitrite (bottom) concentrations. Black horizontal lines indicate target influent dose.

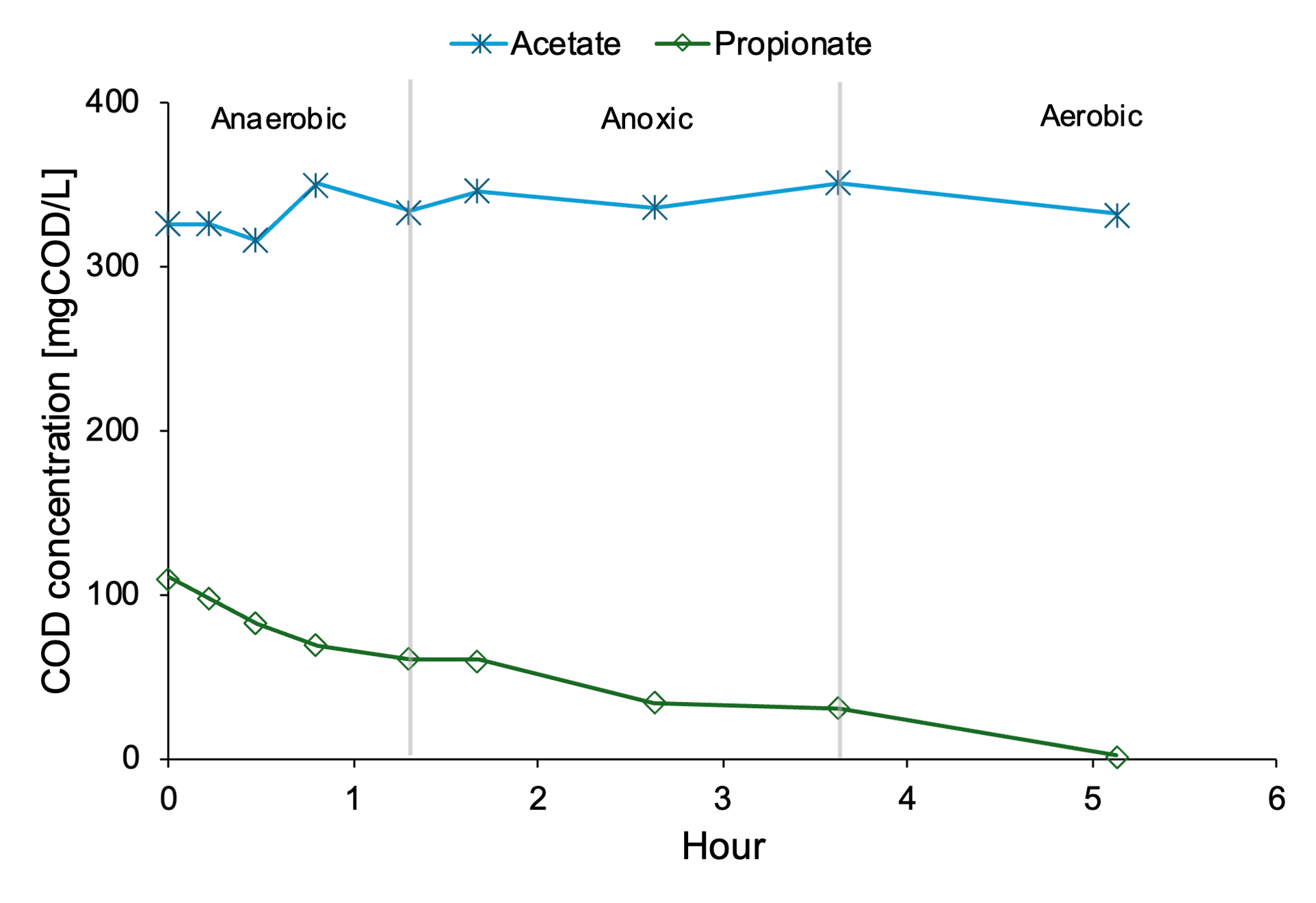

Figure 2 Carbon concentration profiles on day 505 (Phase III).

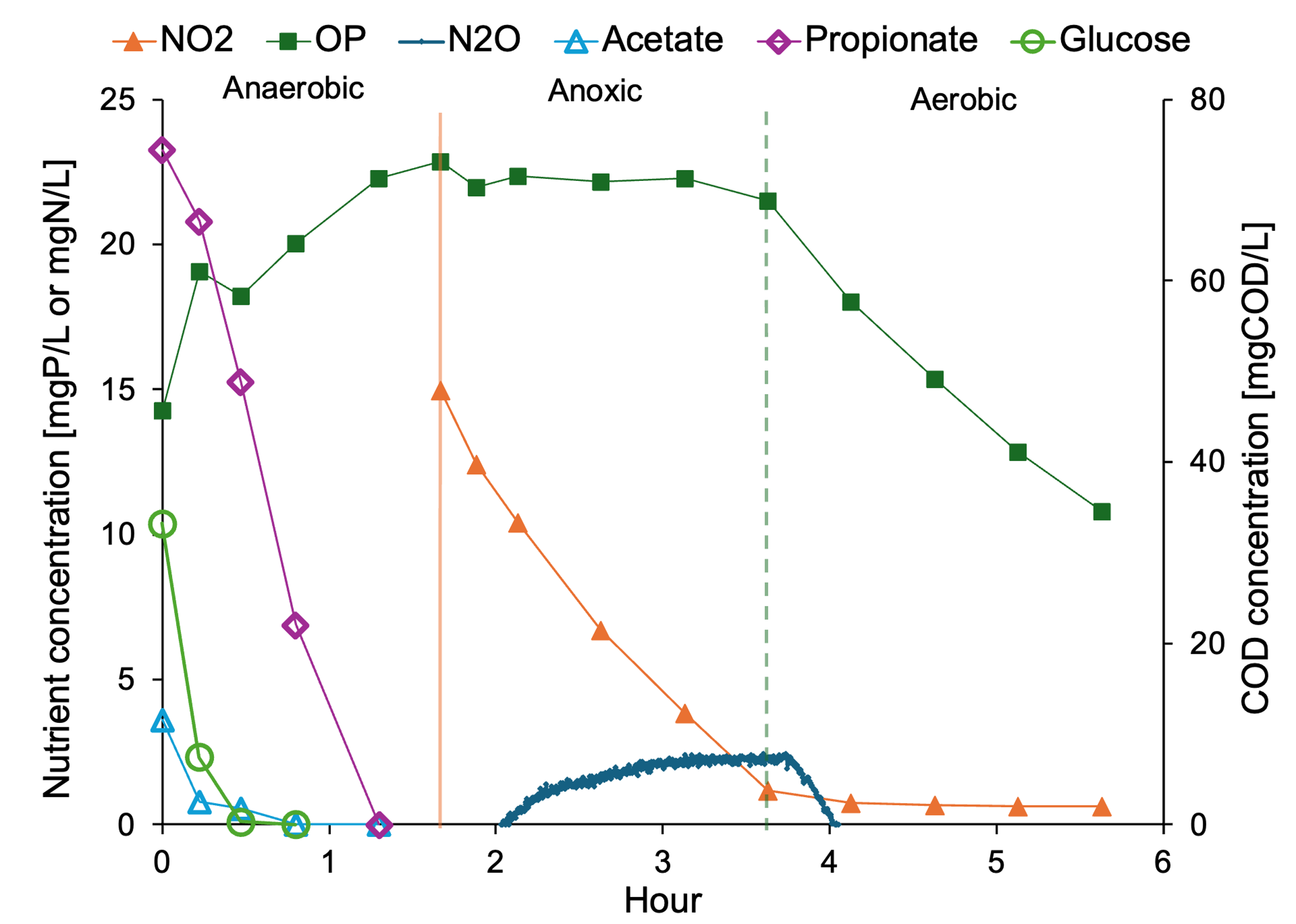

Figure 3 In-cycle nutrient and carbon profile on day 553 (Phase IV).

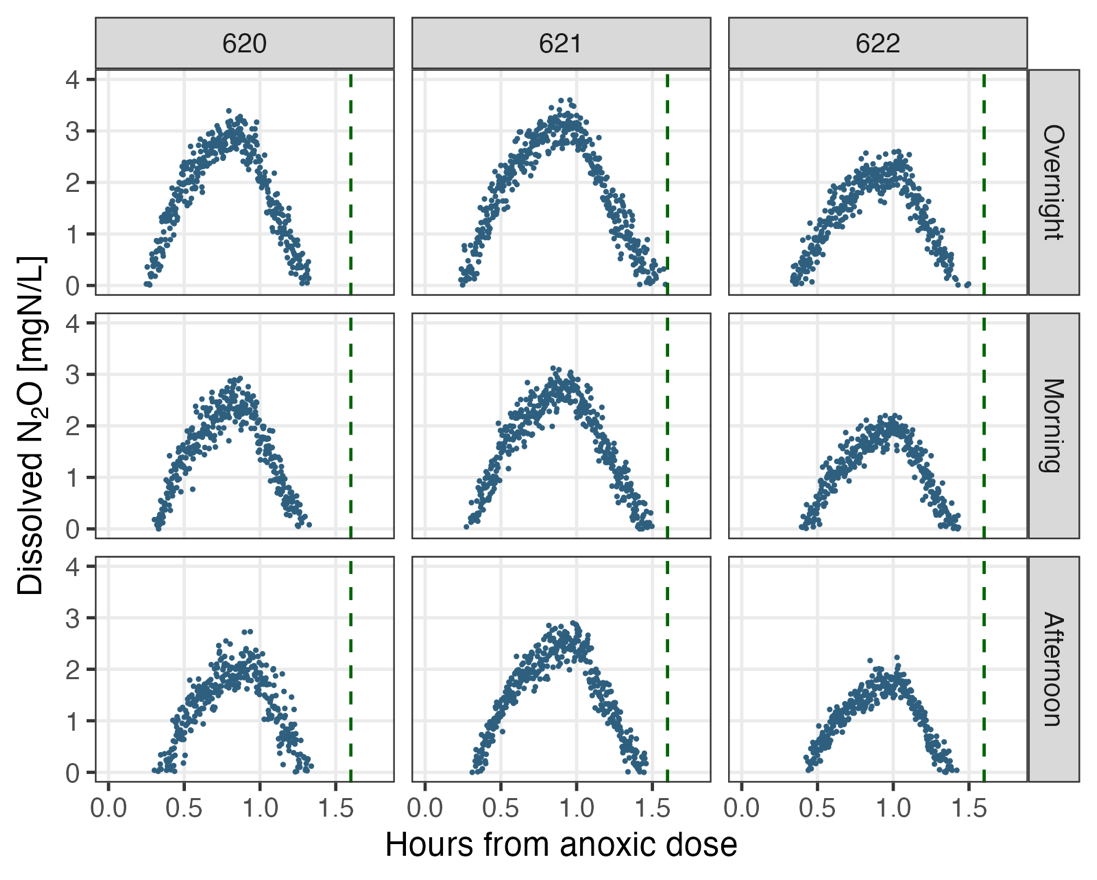

Figure 4 Representative N_2_O profiles during Phase IV. The green vertical line indicates the start of the aerobic phase. The NO_2_ dose was approximately 30 mgN/L during this time period.

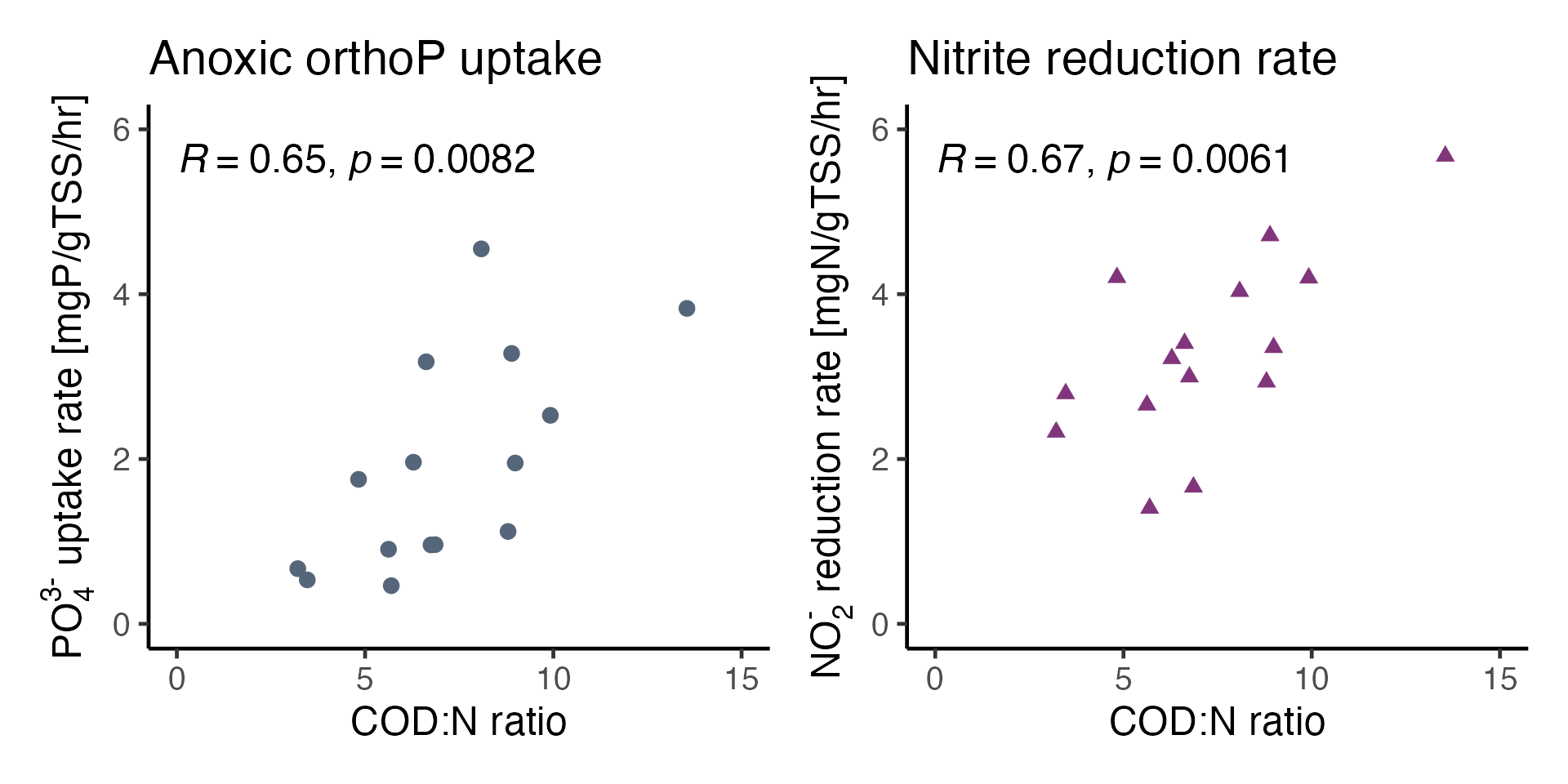

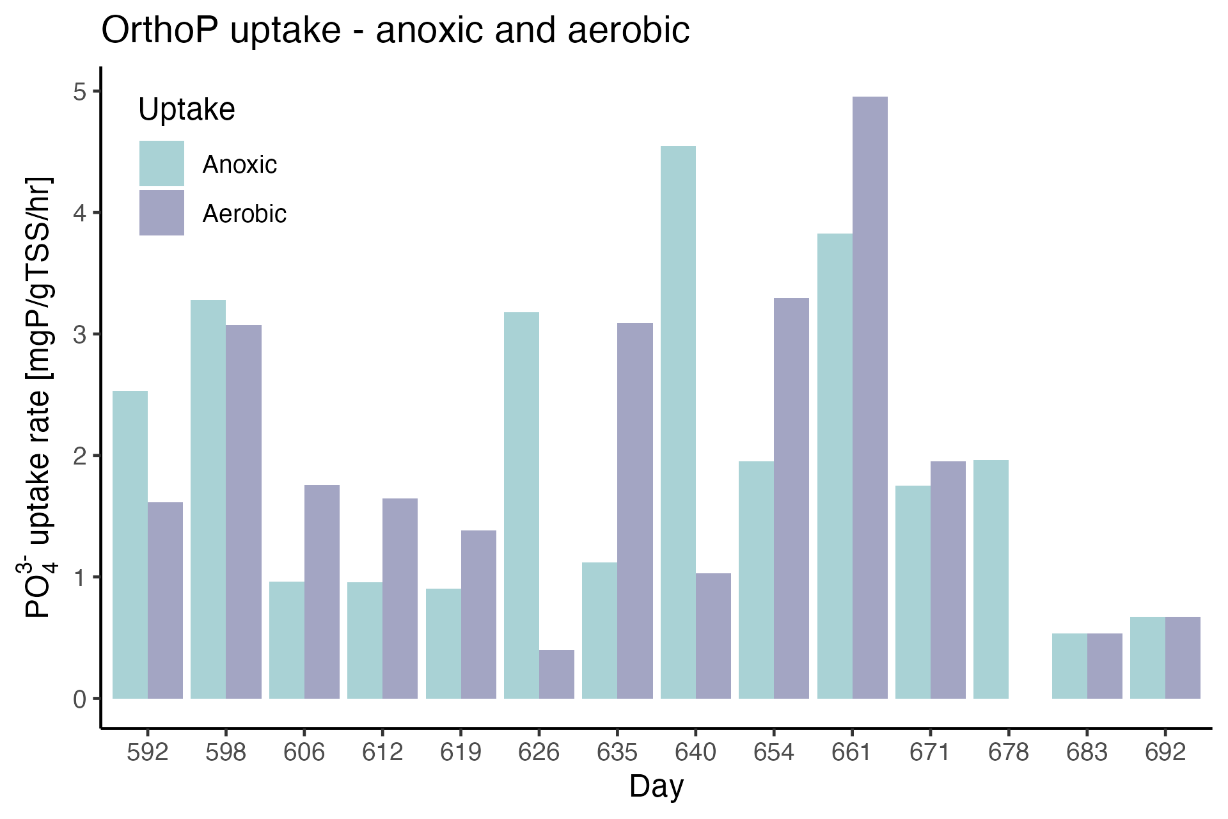

Figure 5 Anoxic orthoP uptake (top left) and NO_2_^-^ reduction rates (top right) plotted against COD:N ratio, and orthoP anoxic and aerobic uptake rates (bottom) measured during in-cycle sampling from Phase IV. Pearson correlation coefficient and significance are provided in the top plots.

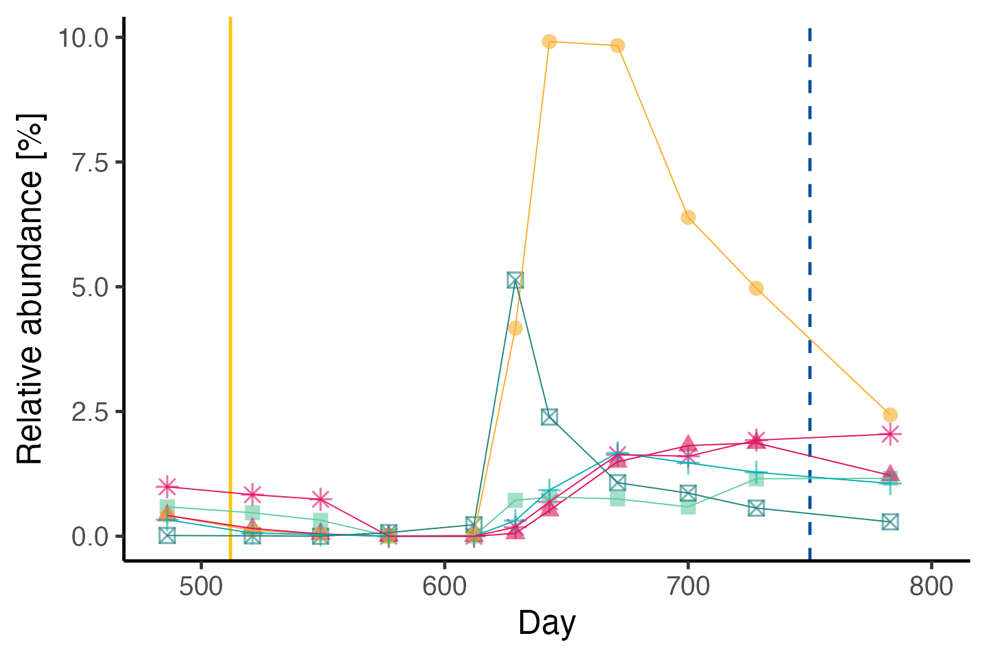

Figure 6 *Ca*. Competibacter amplicon sequence variants (ASVs) during Phases III, IV, and V. The yellow line indicates the start of Phase IV and the dotted blue line indicates the start of Phase V.

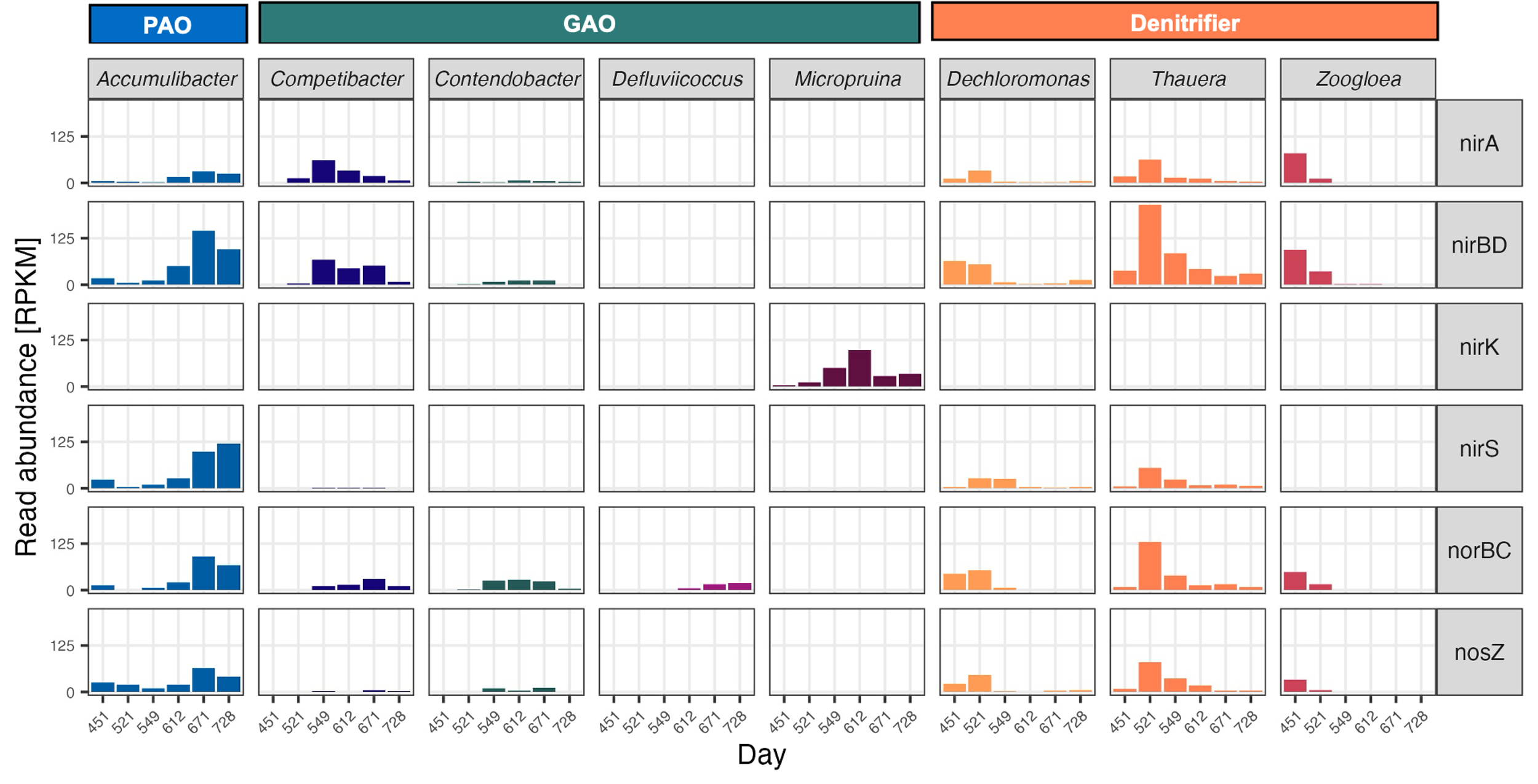

Figure 7 PAO, GAO, and denitrifier gene abundances of denitrification pathway genes based on metagenomic analysis. Day 451 is from Phase II and the rest of the timepoints are from Phase IV.

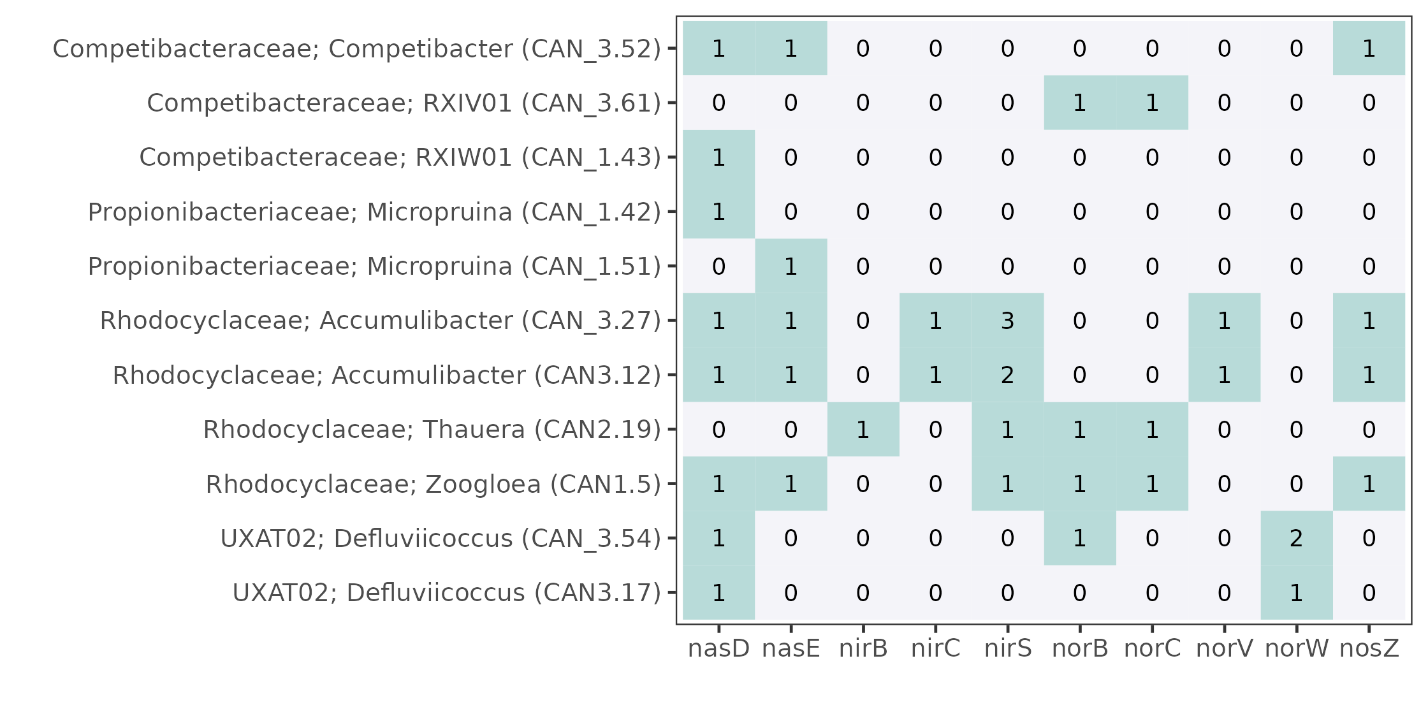

Figure 8 Denitrification pathway gene annotations in PAO, GAO, and denitrifier metagenome-assembled genomes. MAGs are labeled by family, genus and MAG ID. The number labels show copy numbers.

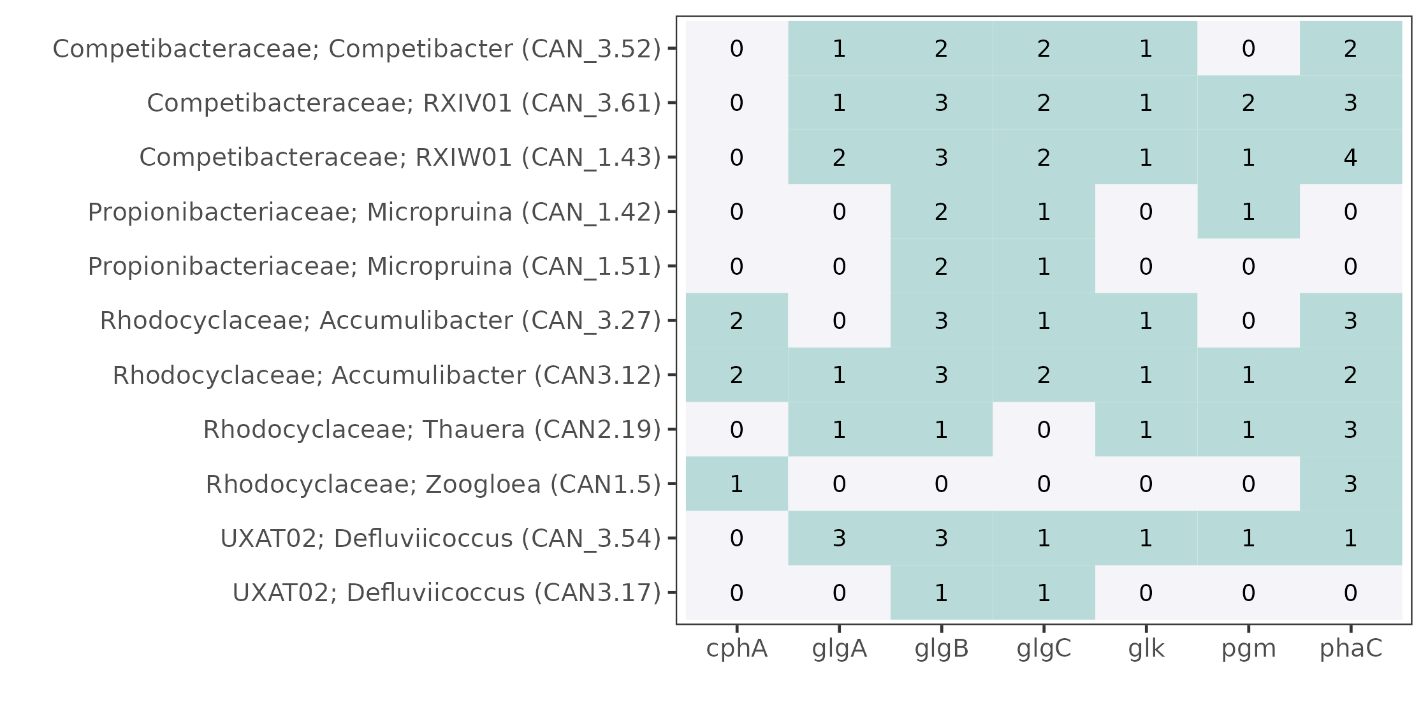

Figure 9 Carbon uptake and utilization pathway gene annotations in PAO, GAO, and denitrifier metagenome-assembled genomes. MAGs are labeled by family, genus and MAG ID. The number labels show copy numbers.

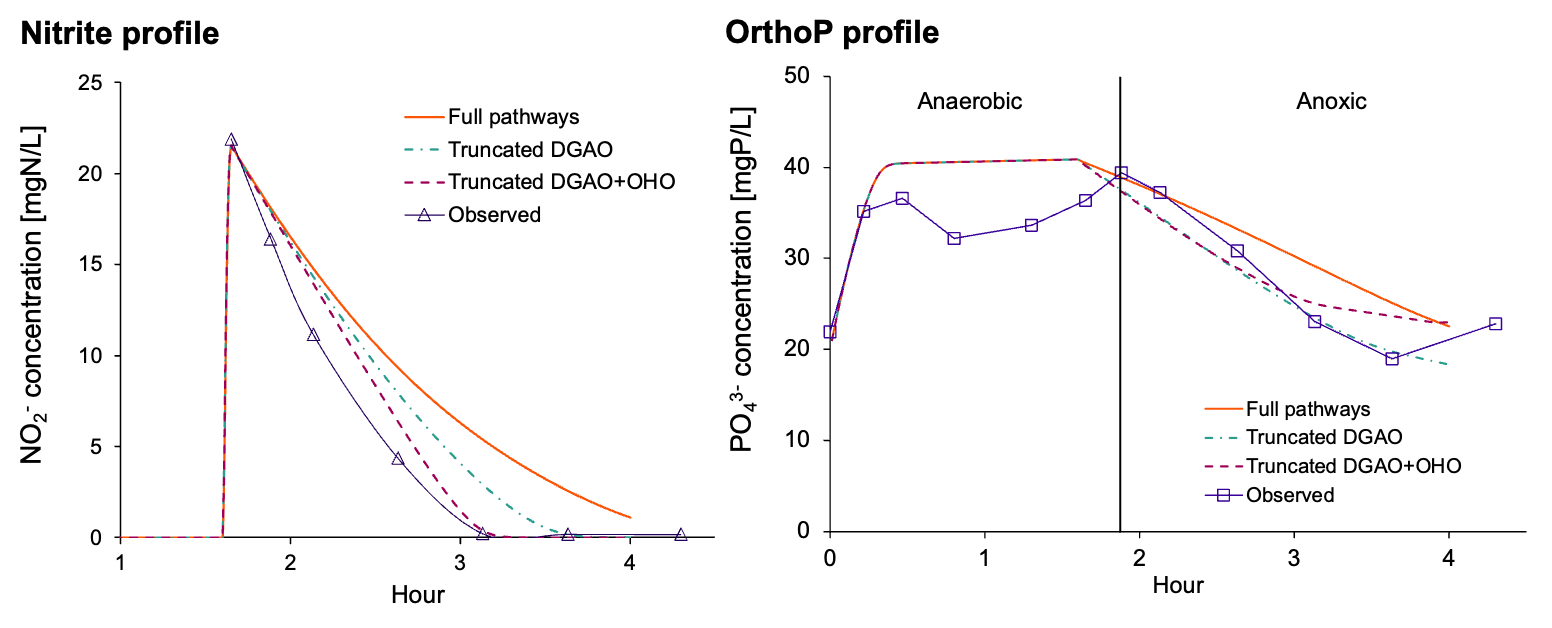

Figure 10 Nirite (left) and orthoP (right) profiles for the modeling scenarios compared to the observed values from day 623.

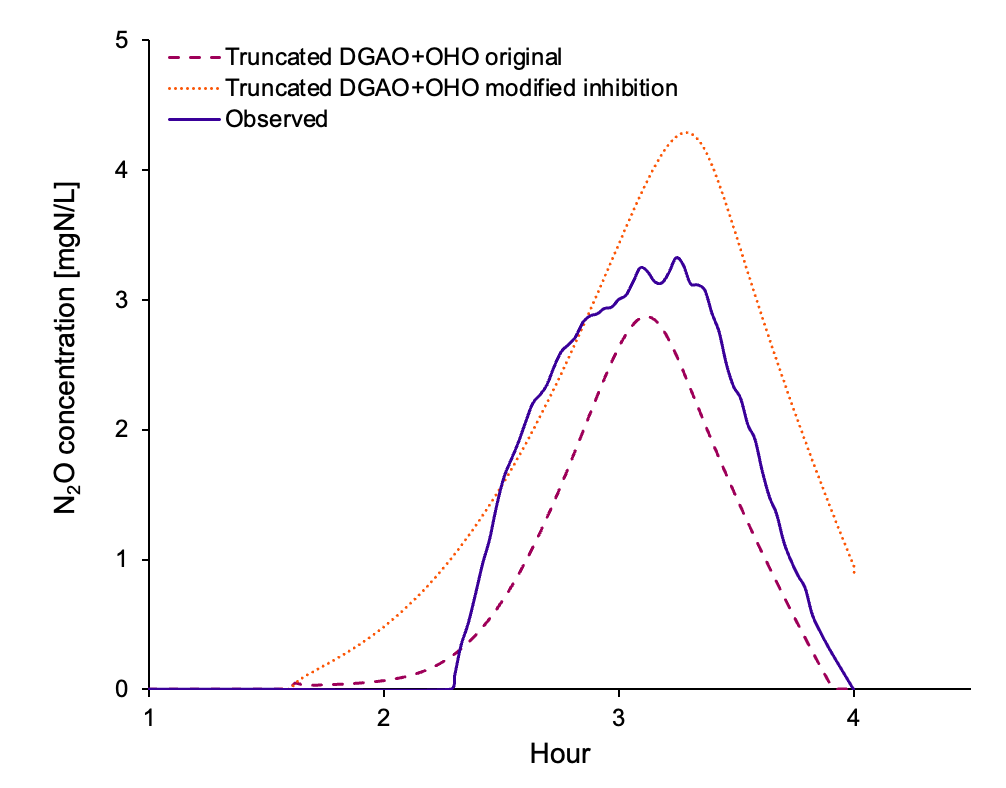

Figure 11 N_2_O profiles of the truncated DGAO+OHO scenario as well as a truncated DGAO+OHO scenario where all NO_2_^-^ inhibition constants were assumed to be the same (8 gN/m^3^).

Table 1 Characteristics of medium and high quality PAO, GAO, and denitrifier metagenome-assembled genomes recovered from this study.

| Day | MAG ID | Classification | Completeness | Contamination |
| --- | --- | --- | --- | --- |
| 451 | CAN1.5 | d__Bacteria;p__Pseudomonadota;c__Gammaproteobacteria;o__Burkholderiales;f__Rhodocyclaceae;g__Zoogloea;s__Zoogloea sp016720685 | 68.83 | 1.29 |
| 521 | CAN2.19 | d__Bacteria;p__Pseudomonadota;c__Gammaproteobacteria;o__Burkholderiales;f__Rhodocyclaceae;g__Thauera;s__Thauera sp016790365 | 74.42 | 1.63 |
| 549 | CAN_1.42 | d__Bacteria;p__Actinomycetota;c__Actinomycetia;o__Propionibacteriales;f__Propionibacteriaceae;g__Micropruina;s__Micropruina glycogenica | 76.95 | 2.54 |
| 549 | CAN_1.43 | d__Bacteria;p__Pseudomonadota;c__Gammaproteobacteria;o__Competibacterales;f__Competibacteraceae;g__RXIW01;s__RXIW01 sp003989065 | 93.94 | 4.3 |
| 549 | CAN_1.51 | d__Bacteria;p__Actinomycetota;c__Actinomycetia;o__Propionibacteriales;f__Propionibacteriaceae;g__Micropruina;s__ | 73.4 | 3.97 |
| 612 | CAN_2.103 | d__Bacteria;p__Actinomycetota;c__Actinomycetia;o__Propionibacteriales;f__Propionibacteriaceae;g__Micropruina;s__ | 65.7 | 4.4 |
| 612 | CAN_2.14 | d__Bacteria;p__Actinomycetota;c__Actinomycetia;o__Actinomycetales;f__Dermatophilaceae;g__Phosphoribacter;s__Phosphoribacter hodrii | 71.88 | 3.15 |
| 612 | CAN_2.16 | d__Bacteria;p__Pseudomonadota;c__Alphaproteobacteria;o__Rhodospirillales_A;f__UXAT02;g__Defluviicoccus;s__ | 75.57 | 1.49 |
| 671 | CAN_3.27 | d__Bacteria;p__Pseudomonadota;c__Gammaproteobacteria;o__Burkholderiales;f__Rhodocyclaceae;g__Accumulibacter;s__Accumulibacter sp000585075 | 74.14 | 1.72 |
| 671 | CAN_3.31 | d__Bacteria;p__Actinomycetota;c__Actinomycetia;o__Propionibacteriales;f__Propionibacteriaceae;g__Micropruina;s__Micropruina glycogenica | 74.2 | 1.34 |
| 671 | CAN_3.52 | d__Bacteria;p__Pseudomonadota;c__Gammaproteobacteria;o__Competibacterales;f__Competibacteraceae;g__Competibacter;s__Competibacter sp014879435 | 82.76 | 1.44 |
| 671 | CAN_3.54 | d__Bacteria;p__Pseudomonadota;c__Alphaproteobacteria;o__Rhodospirillales_A;f__UXAT02;g__Defluviicoccus;s__ | 83.97 | 3.81 |
| 671 | CAN_3.61 | d__Bacteria;p__Pseudomonadota;c__Gammaproteobacteria;o__Competibacterales;f__Competibacteraceae;g__RXIV01;s__RXIV01 sp020846135 | 96.06 | 2.29 |
| 728 | CAN3.12 | d__Bacteria;p__Pseudomonadota;c__Gammaproteobacteria;o__Burkholderiales;f__Rhodocyclaceae;g__Accumulibacter;s__Accumulibacter sp000585075 | 97.95 | 1.98 |
| 728 | CAN3.17 | d__Bacteria;p__Pseudomonadota;c__Alphaproteobacteria;o__Rhodospirillales_A;f__UXAT02;g__Defluviicoccus;s__ | 54.56 | 0.5 |
| 728 | CAN3.18 | d__Bacteria;p__Actinomycetota;c__Actinomycetia;o__Propionibacteriales;f__Propionibacteriaceae;g__Micropruina;s__Micropruina glycogenica | 81.04 | 3.28 |

Table 2 Kinetic rate equations

| Process | Rate expression |
| --- | --- |
| Anaerobic carbon storage by DPAO | $q_{PHA,DPAO}\frac{S_{S}}{K_{S,DPAO}+S_{S}}\frac{X_{\mathrm{PP}}/X_{\mathrm{DPAO}}}{K_{PP,DPAO}+X_{\mathrm{PP}}/X_{\mathrm{DPAO}}}X_{\mathrm{DPAO}}$ |
| Anaerobic carbon storage by DGAO | $q_{PHA, DGAO}\frac{S_{S}}{K_{S,DGAO}+S_{S}}\frac{X_{\mathrm{GLY}}/X_{\mathrm{DGAO}}}{K_{\mathrm{GLY}}+X_{\mathrm{GLY}}/X_{\mathrm{DGAO}}}X_{\mathrm{DGAO}}$ |
| Anaerobic carbon storage by OHO | $q_{STO, OHO}\frac{S_{S}}{K_{S,OHO}+S_{S}}\frac{X_{STO,OHO}/X_{\mathrm{OHO}}}{K_{STO,OHO}+X_{STO, OHO}/X_{\mathrm{OHO}}}X_{\mathrm{OHO}}$ |
| Anoxic storage of X_PP_ on NO_2_^-^ | $q_{\mathrm{PP}}\frac{K_{I,NO2}}{K_{I,NO2}+S_{NO2}}\frac{S_{NO2}}{K_{NO2}+S_{\mathrm{NO}}}\frac{S_{PO4}}{K_{PO4,PP}+S_{PO4}}\frac{X_{PHA,DPAO}/X_{\mathrm{DPAO}}}{K_{PHA,DPAO}+X_{PHA,DPAO}/X_{\mathrm{DPAO}}}\frac{K_{max,DPAO-}X_{\mathrm{PP}}/X_{\mathrm{DPAO}}}{K_{iPP,DPAO}+K_{max,DPAO-}X_{\mathrm{PP}}/X_{\mathrm{DPAO}}}X_{\mathrm{DPAO}}$ |
| Anoxic storage of X_PP_ on NO^-^ | $q_{\mathrm{PP}}\frac{K_{I,NO}}{K_{I,NO}+S_{NO2}}\frac{S_{\mathrm{NO}}}{K_{\mathrm{NO}}+S_{\mathrm{NO}}}\frac{S_{PO4}}{K_{PO4,PP}+S_{PO4}}\frac{X_{PHA,DPAO}/X_{\mathrm{DPAO}}}{K_{PHA,DPAO}+X_{PHA,DPAO}/X_{\mathrm{DPAO}}}\frac{K_{max,DPAO-}X_{\mathrm{PP}}/X_{\mathrm{DPAO}}}{K_{iPP,DPAO}+K_{max,DPAO-}X_{\mathrm{PP}}/X_{\mathrm{DPAO}}}X_{\mathrm{DPAO}}$ |
| Anoxic storage of X_PP_ on N_2_O | $q_{\mathrm{PP}}\frac{K_{I,N2O}}{K_{I,N2O}+S_{NO2}}\frac{S_{N2O}}{K_{N2O}+S_{N2O}}\frac{S_{PO4}}{K_{PO4,PP}+S_{PO4}}\frac{X_{PHA,DPAO}/X_{\mathrm{DPAO}}}{K_{PHA,DPAO}+X_{PHA,DPAO}/X_{\mathrm{DPAO}}}\frac{K_{max,DPAO-}X_{\mathrm{PP}}/X_{\mathrm{DPAO}}}{K_{iPP,DPAO}+K_{max,DPAO-}X_{\mathrm{PP}}/X_{\mathrm{DPAO}}}X_{\mathrm{DPAO}}$ |
| Anoxic storage of X_GLY_ on NO_2_^-^ | $q_{\mathrm{GLY}}\frac{K_{I,NO2}}{K_{I,NO2}+S_{NO2}}\frac{S_{NO2}}{K_{NO2}+S_{\mathrm{NO}}}\frac{X_{PHA,DGAO}/X_{\mathrm{DGAO}}}{K_{PHA,DGAO}+X_{PHA,DGAO}/X_{\mathrm{DGAO}}}\frac{K_{max,DGAO}-X_{\mathrm{GLY}}/X_{\mathrm{DGAO}}}{K_{iGLY,DGAO}+K_{max,DGAO}-X_{\mathrm{GLY}}/X_{\mathrm{DGAO}}}X_{\mathrm{DGAO}}$ |
| Anoxic storage of X_GLY_ on NO^-^ | $q_{\mathrm{GLY}}\frac{K_{I,NO}}{K_{I,NO}+S_{NO2}}\frac{S_{\mathrm{NO}}}{K_{\mathrm{NO}}+S_{\mathrm{NO}}}\frac{X_{PHA,DGAO}/X_{\mathrm{DGAO}}}{K_{PHA,DGAO}+X_{PHA,DGAO}/X_{\mathrm{DGAO}}}\frac{K_{max,DGAO}-X_{\mathrm{GLY}}/X_{\mathrm{DGAO}}}{K_{iGLY,DGAO}+K_{max,DGAO}-X_{\mathrm{GLY}}/X_{\mathrm{DGAO}}}X_{\mathrm{DGAO}}$ |
| Anoxic storage of X_GLY_ on N_2_O | $q_{\mathrm{GLY}}\frac{K_{I,N2O}}{K_{I,N2O}+S_{NO2}}\frac{S_{N2O}}{K_{N2O}+S_{N2O}}\frac{X_{PHA,DGAO}/X_{\mathrm{DGAO}}}{K_{PHA,DGAO}+X_{PHA,DGAO}/X_{\mathrm{DGAO}}}\frac{K_{max,DGAO}-X_{\mathrm{GLY}}/X_{\mathrm{DGAO}}}{K_{iGLY,DGAO}+K_{max,DGAO}-X_{\mathrm{GLY}}/X_{\mathrm{DGAO}}}X_{\mathrm{DGAO}}$ |
| Anoxic growth on NO_2_^-^by DPAO | $\mu_{DPAO,NO2}\frac{K_{I,NO2}}{K_{I,NO2}+S_{NO2}}\frac{S_{NO2}}{K_{NO2}+S_{NO2}}\frac{S_{PO4}}{K_{DPAO,PO4}+S_{PO4}}\frac{{X_{PHA,DPAO}}/{X_{\mathrm{DPAO}}}}{K_{PHA,DPAO}+{X_{PHA,DPAO}}/{X_{\mathrm{DPAO}}}}X_{\mathrm{DPAO}}$ |
| Anoxic growth on NO by DPAO | $\mu_{DPAO,NO}\frac{K_{I,NO}}{K_{I,NO}+S_{NO2}}\frac{S_{\mathrm{NO}}}{K_{\mathrm{NO}}+S_{\mathrm{NO}}}\frac{S_{PO4}}{K_{DPAO,PO4}+S_{PO4}}\frac{{X_{PHA,DPAO}}/{X_{\mathrm{DPAO}}}}{K_{PHA,DPAO}+{X_{PHA,DPAO}}/{X_{\mathrm{DPAO}}}}X_{\mathrm{DPAO}}$ |
| Anoxic growth on N_2_O by DPAO | $\mu_{DPAO,N2O}\frac{K_{I,N2O}}{K_{I,N2O}+S_{NO2}}\frac{S_{N2O}}{K_{N2O}+S_{N2O}}\frac{S_{PO4}}{K_{DPAO,PO4}+S_{PO4}}\frac{{X_{PHA,DPAO}}/{X_{\mathrm{DPAO}}}}{K_{PHA,DPAO}+{X_{PHA,DPAO}}/{X_{\mathrm{DPAO}}}}X_{\mathrm{DPAO}}$ |
| Anoxic growth on NO_2_^-^by DGAO | $\mu_{DGAO,NO2}\frac{K_{I,NO2}}{K_{I,NO2}+S_{NO2}}\frac{S_{NO2}}{K_{NO2}+S_{NO2}}\frac{{X_{PHA,DGAO}}/{X_{\mathrm{DGAO}}}}{K_{PHA,DGAO}+{X_{PHA,DGAO}}/{X_{\mathrm{DGAO}}}}X_{\mathrm{DGAO}}$ |
| Anoxic growth on NO by DGAO | $\mu_{DGAO,NO}\frac{K_{I,NO}}{K_{I,NO}+S_{NO2}}\frac{S_{\mathrm{NO}}}{K_{\mathrm{NO}}+S_{\mathrm{NO}}}\frac{{X_{PHA,DGAO}}/{X_{\mathrm{DGAO}}}}{K_{PHA,DGAO}+{X_{PHA,DGAO}}/{X_{\mathrm{DGAO}}}}X_{\mathrm{DGAO}}$ |
| Anoxic growth on N_2_O by DGAO | $\mu_{DGAO,N2O}\frac{K_{I,N2O}}{K_{I,N2O}+S_{NO2}}\frac{S_{N2O}}{K_{N2O}+S_{N2O}}\frac{{X_{PHA,DGAO}}/{X_{\mathrm{DGAO}}}}{K_{PHA,DGAO}+{X_{PHA,DGAO}}/{X_{\mathrm{DGAO}}}}X_{\mathrm{DGAO}}$ |
| Anoxic growth on NO_2_^-^by OHO | $\mu_{OHO,NO2}\frac{K_{I,NO2}}{K_{I,NO2}+S_{NO2}}\frac{S_{NO2}}{K_{NO2}+S_{NO2}}\frac{{X_{STO,OHO}}/{X_{\mathrm{OHO}}}}{K_{STO, OHO}+{X_{STO,OHO}}/{X_{\mathrm{OHO}}}}X_{\mathrm{OHO}}$ |
| Anoxic growth on NO by OHO | $\mu_{OHO,NO}\frac{K_{I,NO}}{K_{I,NO}+S_{NO2}}\frac{S_{\mathrm{NO}}}{K_{\mathrm{NO}}+S_{\mathrm{NO}}}\frac{{X_{STO,OHO}}/{X_{\mathrm{OHO}}}}{K_{STO, OHO}+{X_{STO,OHO}}/{X_{\mathrm{OHO}}}}X_{\mathrm{OHO}}$ |
| Anoxic growth on N_2_O by OHO | $\mu_{OHO,N2O}\frac{K_{I,N2O}}{K_{I,N2O}+S_{NO2}}\frac{S_{N2O}}{K_{N2O}+S_{N2O}}\frac{{X_{STO,OHO}}/{X_{\mathrm{OHO}}}}{K_{STO, OHO}+{X_{STO,OHO}}/{X_{\mathrm{OHO}}}}X_{\mathrm{OHO}}$ |
| Decay of DPAO | $b_{\mathrm{DPAO}}X_{\mathrm{DPAO}}$ |
| Decay of X_PP_ | $b_{\mathrm{PP}}X_{\mathrm{PP}}$ |
| Decay of X_PHA,DPAO_ | $b_{PHA,DPAO}X_{PHA,DPAO}$ |
| Decay of DGAO | $b_{\mathrm{DGAO}}X_{\mathrm{DGAO}}$ |
| Decay of X_GLY_ | $b_{\mathrm{GLY}}X_{\mathrm{GLY}}$ |
| Decay of X_PHA,DGAO_ | $b_{PHA,DGAO}X_{PHA,DGAO}$ |
| Decay of X_OHO_ | $b_{\mathrm{OHO}}X_{\mathrm{OHO}}$ |
| Decay of X_STO_ | $b_{\mathrm{STO}}X_{STO,OHO}$ |

Table 3 Stoichiometric matrix

| Process | S_NO2_ | S_NO_ | S_N2O_ | S_N2_ | S_S_ | S_PO4_ | X_DPAO_ | X_DGAO_ | X_OHO_ | X_PP_ | X_GLY_ | X_PHA,DPAO_ | X_PHA,DGAO_ | X_STO,OHO_ | X_I_ |
| --- | --- | --- | --- | --- | --- | --- | --- | --- | --- | --- | --- | --- | --- | --- | --- |
| Anaerobic carbon storage by DPAO |  |  |  |  | -1 | Y_PO4_ |  |  |  | -Y_PO4_ |  | 1 |  |  |  |
| Anaerobic carbon storage by DGAO |  |  |  |  | -1 |  |  |  |  |  | -Y_GLY_ |  | 1 |  |  |
| Anaerobic carbon storage by OHO |  |  |  |  | -1 |  |  |  |  |  |  |  |  | Y_STO_ |  |
| Anoxic storage of X_PP_ on NO_2_^-^ | $-\frac{Y_{\mathrm{PHA}}}{0.57}$ | $\frac{Y_{\mathrm{PHA}}}{0.57}$ |  |  |  | -1 |  |  |  | 1 |  | -Y_PHA_ |  |  |  |
| Anoxic storage of X_PP_ on NO^-^ |  | $-\frac{Y_{\mathrm{PHA}}}{0.57}$ | $\frac{Y_{\mathrm{PHA}}}{0.57}$ |  |  | -1 |  |  |  | 1 |  | -Y_PHA_ |  |  |  |
| Anoxic storage of X_PP_ on N_2_O |  |  | $-\frac{Y_{\mathrm{PHA}}}{0.57}$ | $\frac{Y_{\mathrm{PHA}}}{0.57}$ |  | -1 |  |  |  | 1 |  | -Y_PHA_ |  |  |  |
| Anoxic storage of X_GLY_ on NO_2_^-^ | $-\frac{Y_{\mathrm{GLY}}}{0.57}$ | $\frac{Y_{\mathrm{GLY}}}{0.57}$ |  |  |  |  |  |  |  |  | 1 |  | -Y_GLY_ |  |  |
| Anoxic storage of X_GLY_ on NO^-^ |  | $-\frac{Y_{\mathrm{GLY}}}{0.57}$ | $\frac{Y_{\mathrm{GLY}}}{0.57}$ |  |  |  |  |  |  |  | 1 |  | -Y_GLY_ |  |  |
| Anoxic storage of X_GLY_ on N_2_O |  |  | $-\frac{Y_{\mathrm{GLY}}}{0.57}$ | $\frac{Y_{\mathrm{GLY}}}{0.57}$ |  |  |  |  |  |  | 1 |  | -Y_GLY_ |  |  |
| Anoxic growth on NO_2_^-^by DPAO | $-\frac{{1-Y}_{DPAO,NOx}}{0.57Y_{DPAO,NOx}}$ | $\frac{{1-Y}_{DPAO,NOx}}{0.57Y_{DPAO,NOx}}$ |  |  |  | -i_P,BM_ | 1 |  |  |  |  | $-\frac{1}{Y_{DPAO,NOx}}$ |  |  |  |
| Anoxic growth on NO by DPAO |  | $-\frac{{1-Y}_{DPAO,NOx}}{0.57Y_{DPAO,NOx}}$ | $\frac{{1-Y}_{DPAO,NOx}}{0.57Y_{DPAO,NOx}}$ |  |  | -i_P,BM_ | 1 |  |  |  |  | $-\frac{1}{Y_{DPAO,NOx}}$ |  |  |  |
| Anoxic growth on N_2_O by DPAO |  |  | $-\frac{{1-Y}_{DPAO,NOx}}{0.57Y_{DPAO,NOx}}$ | $\frac{{1-Y}_{DPAO,NOx}}{0.57Y_{DPAO,NOx}}$ |  | -i_P,BM_ | 1 |  |  |  |  | $-\frac{1}{Y_{DPAO,NOx}}$ |  |  |  |
| Anoxic growth on NO_2_^-^by DGAO | $-\frac{{1-Y}_{DGAO,NOx}}{0.57Y_{DGAO,NOx}}$ | $\frac{{1-Y}_{DGAO,NOx}}{0.57Y_{DGAO,NOx}}$ |  |  |  | -i_P,BM_ | 1 |  |  |  |  |  | $-\frac{1}{Y_{DGAO,NOx}}$ |  |  |
| Anoxic growth on NO by DGAO |  | $-\frac{{1-Y}_{DGAO,NOx}}{0.57Y_{DGAO,NOx}}$ | $\frac{{1-Y}_{DGAO,NOx}}{0.57Y_{DGAO,NOx}}$ |  |  | -i_P,BM_ | 1 |  |  |  |  |  | $-\frac{1}{Y_{DGAO,NOx}}$ |  |  |
| Anoxic growth on N_2_O by DGAO |  |  | $-\frac{{1-Y}_{DGAO,NOx}}{0.57Y_{DGAO,NOx}}$ | $\frac{{1-Y}_{DGAO,NOx}}{0.57Y_{DGAO,NOx}}$ |  | -i_P,BM_ | 1 |  |  |  |  |  | $-\frac{1}{Y_{DGAO,NOx}}$ |  |  |
| Anoxic growth on NO_2_^-^by OHO | $-\frac{{1-Y}_{OHO,NOx}}{0.57Y_{OHO,NOx}}$ | $\frac{{1-Y}_{OHO,NOx}}{0.57Y_{OHO,NOx}}$ |  |  |  | -i_P,BM_ | 1 |  |  |  |  |  |  | $-\frac{1}{Y_{OHO,NOx}}$ |  |
| Anoxic growth on NO by OHO |  | $-\frac{{1-Y}_{OHO,NOx}}{0.57Y_{OHO,NOx}}$ | $\frac{{1-Y}_{OHO,NOx}}{0.57Y_{OHO,NOx}}$ |  |  | -i_P,BM_ | 1 |  |  |  |  |  |  | $-\frac{1}{Y_{OHO,NOx}}$ |  |
| Anoxic growth on N_2_O by OHO |  |  | $-\frac{{1-Y}_{OHO,NOx}}{0.57Y_{OHO,NOx}}$ | $\frac{{1-Y}_{OHO,NOx}}{0.57Y_{OHO,NOx}}$ |  | -i_P,BM_ | 1 |  |  |  |  |  |  | $-\frac{1}{Y_{OHO,NOx}}$ |  |
| Decay of DPAO |  |  |  |  |  | $i_{P,BM}-i_{P,XI}f_{I}$ | -1 |  |  |  |  |  |  |  | $f_{I}$ |
| Decay of X_PP_ |  |  |  |  |  | 1 |  |  |  | -1 |  |  |  |  |  |
| Decay of X_PHA,DPAO_ |  |  |  |  |  |  |  |  |  |  |  | -1 |  |  |  |
| Decay of DGAO |  |  |  |  |  | $i_{P,BM}-i_{P,XI}f_{I}$ |  | -1 |  |  |  |  |  |  | $f_{I}$ |
| Decay of X_GLY_ |  |  |  |  |  |  |  |  |  |  | -1 |  |  |  |  |
| Decay of X_PHA,DGAO_ |  |  |  |  |  |  |  |  |  |  |  |  | -1 |  |  |
| Decay of X_OHO_ |  |  |  |  |  | $i_{P,BM}-i_{P,XI}f_{I}$ |  |  | -1 |  |  |  |  |  | $f_{I}$ |
| Decay of X_STO_ |  |  |  |  |  |  |  |  |  |  |  |  |  | -1 |  |

Table 4 Stoichiometric and kinetic parameters

| Parameter | Definition | Value | Unit | Source |
| --- | --- | --- | --- | --- |
| Stochiometric | |  |  |  |
| Y_PO4_ | X_PP_ required per X_PHA,DPAO_ stored | 0.35 | gP/gCOD | (Rieger et al., 2001) |
| Y_PHA_ | X_PHA_ required per X_PP_ stored | 0.2 | gCOD/gP | (Rieger et al., 2001) |
| Y_GLY_ | X_GLY_ required per X_PHA,DGAO_ stored | 0.5 | gCOD/gCOD | (Ren et al., 2023) |
| Y_STO_ | X_STO_ produced per S_S_ | 0.7 | gCOD/gCOD | (Rieger et al., 2001) |
| Y_DPAO,NOx_ | Anoxic yield coefficient for X_DPAO_ | 0.5 | gCOD/gCOD | (Rieger et al., 2001) |
| Y_DGAO,NOx_ | Anoxic yield coefficient for X_DGAO_ | 0.5 | gCOD/gCOD | (Rieger et al., 2001) |
| Y_OHO,NOx_ | Anoxic yield coefficient for X_OHO_ | 0.5 | gCOD/gCOD | (Rieger et al., 2001) |
| i_P,BM_ | Phosphorus content of biomass | 0.02 | gP/gCOD | (Rieger et al., 2001) |
| i_P,XI_ | Phosphorus content of X_I_ | 0.01 | gP/gCOD | (Rieger et al., 2001) |
| f_I_ | Production of X_I_ from endogenous respiration | 0.2 | gCOD/gCOD | (Rieger et al., 2001) |
| Kinetic | |  |  |  |
| q_PHA,DPAO_ | Rate constant for X_PHA_ storage by DPAO | 0.53 | 1/h | (Liu et al., 2015a) |
| q_PHA,DGAO_ | Rate constant for X_PHA_ storage by DGAO | 0.89 | 1/h | (Ren et al., 2023) |
| q_STO,OHO_ | Rate constant for X_STO_ storage by OHO | 0.89 | 1/h | (Ren et al., 2023) |
| q_PP_ | Rate constant for X_PP_ storage | 0.0625 | 1/h | (Rieger et al., 2001) |
| q_GLY_ | Rate constant for X_GLY_ storage | 0.23 | 1/h | (Ren et al., 2023) |
| K_I,NO2_ | Inhibition constant of NO_2_ on NO_2_ reduction | 10 | gN/m3 | (Liu et al., 2015b) |
| K_I,NO_ | Inhibition constant of NO_2_ on NO reduction | 17 | gN/m3 | (Liu et al., 2015b) |
| K_I,N2O_ | Inhibition constant of NO_2_ on N_2_O reduction | 8 | gN/m3 | (Liu et al., 2015b) |
| K_S,DPAO_ | Saturation constant for DPAO on S_S_ | 10 | gCOD/m^3^ | (Rieger et al., 2001) |
| K_S,DGAO_ | Saturation constant for DGAO on S_S_ | 10 | gCOD/m^3^ | (Rieger et al., 2001) |
| K_S,OHO_ | Saturation constant for OHO on S_S_ | 10 | gCOD/m^3^ | (Rieger et al., 2001) |
| K_PP,DPAO_ | Saturation constant for X_PP_/X_DPAO_ | 0.05 | gP/gCOD | (Rieger et al., 2001) |
| K_PO4,PP_ | Saturation constant of S_PO4_ for X_PP_ storage | 0.2 | gP/m^3^ | (Rieger et al., 2001) |
| K_PHA,DPAO_ | Saturation constant for DPAO on X_PHA_ | 0.1 | gCOD/gCOD | (Rieger et al., 2001) |
| K_max,DPAO_ | Maximum ratio of X_PP_/X_DPAO_ | 0.2 | gP/gCOD | (Rieger et al., 2001) |
| K_iPP,DPAO_ | Saturation constant for K_max,DPAO_ –X_PP_/X_DPAO_ | 0.05 | gP/gCOD | (Rieger et al., 2001) |
| K_PHA,DGAO_ | Saturation constant for DGAO on X_PHA_ | 0.1 | gCOD/gCOD | (Ren et al., 2023) |
| K_max,DGAO_ | Maximum ratio of X_GLY_/X_DGAO_ | 0.2 | gCOD/gCOD | (Ren et al., 2023) |
| K_iGLY,DGAO_ | Saturation constant for K_max,DGAO_ – X_GLY_/X_DPAO_ | 0.05 | gCOD/gCOD | (Ren et al., 2023) |
| K_STO,OHO_ | Saturation constant for OHO on X_STO_ | 0.1 | gCOD/gCOD | (Rieger et al., 2001) |
| µ_DPAO,NO2_ | Anoxic growth rate of DPAO on nitrite | 0.019 | 1/h | (Liu et al., 2015a) |
| µ_DPAO,NO_ | Anoxic growth rate of DPAO on nitric oxide | 0.142 | 1/h | (Ni et al., 2011) |
| µ_DPAO,N2O_ | Anoxic growth rate of DPAO on nitrous oxide | 0.018 | 1/h | (Liu et al., 2015a) |
| µ_DGAO,NO2_ | Anoxic growth rate of DGAO on nitrite | 0.021 | 1/h | (Ren et al., 2023) |
| µ_DGAO,NO_ | Anoxic growth rate of DGAO on nitric oxide | 0.091 | 1/h | (Ren et al., 2023) |
| µ_DGAO,N2O_ | Anoxic growth rate of DGAO on nitrous oxide | 0.032 | 1/h | (Ren et al., 2023) |
| µ_OHO,NO2_ | Anoxic growth rate of OHO on nitrite | 0.056 | 1/h | (Ni et al., 2011) |
| µ_OHO,NO_ | Anoxic growth rate of OHO on nitric oxide | 0.142 | 1/h | (Ni et al., 2011) |
| µ_OHO,N2O_ | Anoxic growth rate of OHO on nitrous oxide | 0.134 | 1/h | (Ni et al., 2011) |
| K_NO2_ | S_NO2_ affinity constant | 0.81 | gN/m^3^ | (Ni et al., 2011) |
| K_NO_ | S_NO_ affinity constant | 0.0021 | gN/m^3^ | (Ni et al., 2011) |
| K_N2O_ | S_N2O_ affinity constant | 0.0052 | gN/m^3^ | (Ni et al., 2011) |
| K_NOx_ | Biomass S_NOx_ affinity constant | 0.5 | gN/m^3^ | (Rieger et al., 2001) |
| b_DPAO_ | Decay rate of X_DPAO_ | 0.005 | 1/h | (Rieger et al., 2001) |
| b_PP_ | Decay rate of X_PP_ | 0.005 | 1/h | (Rieger et al., 2001) |
| b_PHA,DPAO_ | Decay rate of X_PHA,DPAO_ by DPAO | 0.005 | 1/h | (Rieger et al., 2001) |
| b_DGAO_ | Decay rate of X_DGAO_ | 0.005 | 1/h | (Rieger et al., 2001) |
| b_GLY_ | Decay rate of X_GLY_ by DGAO | 0.005 | 1/h | (Rieger et al., 2001) |
| b_PHA,DGAO_ | Decay rate of X_PHA,DGAO_ by DGAO | 0.005 | 1/h | (Rieger et al., 2001) |
| b_OHO_ | Decay rate of X_OHO_ | 0.026 | 1/h | (Liu et al., 2015b) |
| b_STO_ | Decay rate of X_STO_ by OHO | 0.004 | 1/h | (Rieger et al., 2001) |
